# A tomato ethylene-insensitive mutant displays altered growth and higher β-carotene levels in fruit

**DOI:** 10.1101/2023.11.14.566984

**Authors:** Suresh Kumar Gupta, Parankusam Santisree, Prateek Gupta, Himabindu Vasuki Kilambi, Yellamaraju Sreelakshmi, Rameshwar Sharma

**Author notes:** **E-mails authors:** (SKG), (PS), (PG), (YS), (RS).

## Abstract

The mutants insensitive to ethylene are helpful in deciphering the role of ethylene in plant development. We isolated an ethylene-insensitive tomato (*Solanum lycopersicum*) mutant by screening for acetylene-resistant (*atr-1*) seedlings. The *atr-1* mutant displayed resistance to kinetin, suggesting attenuation of the ethylene sensing response. *atr-1* also exhibited resistance to ABA- and glucose-mediated inhibition of seed germination. Unlike the *Never-ripe* (*Nr*) mutant, *atr-1* seedlings were resistant to glucose, indicating ethylene sensing in *atr-1* is located in a component distinct from *Nr*. Metabolically, *atr-1* seedlings had lower levels of amino acids but higher levels of several phytohormones, including ABA. *atr-1* plants grew faster and produced more flowers, leading to a higher fruit set. However, the *atr- 1* fruits took a longer duration to reach the red-ripe (RR) stage. The ripened *atr-1* fruits had higher β-carotene levels, retained high β-carotene and lycopene levels post-RR stage. The metabolome profiles of post-RR stage *atr-1* fruits revealed increased levels of sugars. The *atr-1* had a P279L mutation in the GAF domain of the *ETR4*, a key ethylene receptor regulating tomato ripening. Our study highlights that novel alleles in ethylene receptors may aid in enhancing the nutritional quality of tomato.

## Introduction

Ethylene, a gaseous hormone, plays multifaceted roles in plant development, including seedling growth, organ senescence, abscission, and ripening of climacteric fruits (**Abeles et al., 1992**). Most knowledge about ethylene biosynthesis and emanating signaling pathways has been obtained by physiological and biochemical characterization of the mutants obtained in different plant species. The above studies revealed that ethylene is perceived at the endoplasmic reticulum (ER) by a family of ER-bound ethylene receptors (ETRs), which inactivates the CTR1 protein kinase, which in turn allows activation of EIN2, a central player in the ethylene-signaling pathway. The activation of EIN2 influences the activity of transcription factors such as EIN3 and EILs, which move to the nucleus and regulate the expression of ethylene-responsive genes (**Binder, 2020**).

Ethylene receptors (ETRs) have homology to bacterial two-component receptors and are conserved across the plant kingdom (**Schaller et al., 2011**). ETRs are composed of several domains; the N-terminal transmembrane domain contains an ethylene-binding site, followed by a cytosolic GAF domain and a kinase domain. Some ETR family members at the C-terminus have a receiver domain (**Binder, 2008**). The ETRs act as negative regulators of the ethylene-signaling pathway, following an inverse-agonist model where ethylene binding turns off their function (**Hall et al., 1999**).

Tomato, being a climacteric fruit, has been used as a model system for studying the role of ethylene in regulating fruit ripening (**Klee and Giovannoni, 2011**). The inhibition of either ethylene production (**Oeller et al., 1991**) or perception leads to improper ripening of tomato fruits (**Wilkinson et al., 1995**). In tomato, at least seven *ETRs* (*LeETR1-7*) have been identified (**Liu et al., 2015**). Tomato ETRs fall into two subfamilies with SlETR1, SlETR2, and SlETR3 (aka NR- Never ripe) in subfamily I and SlETR4, SlETR5, SlETR6, and SlETR7 in subfamily II (**Chen et al., 2018**).

ETRs function as homo-or heteromers, and homodimers may form higher-order clusters with GAF-GAF interaction with different ETRs (**Gao et al., 2008**). These heteromeric complexes possibly fine-tune the signal output from the receptors (**Shakeel et al., 2013**). The formation of such heteromeric complexes has also been observed during tomato fruit ripening using a coimmunoprecipitation (co-IP) assay. The SlETR4 preferentially forms heteromeric receptor complexes with the members of subfamily I (SlETR1, SlETR2, and SlETR3) but not with members of subfamily II (SlETR5, SlETR6, and SlETR7). (**Kamiyoshihara et al., 2022**). SlETR4 also shows co-IP with SlCTR1 and SlCTR3, indicating that SlETR4 forms a wide range of complexes, including downstream signaling components.

The relative function of these ETRs and protein domains in regulating tomato ripening is being elucidated using mutant and transgenic plants. The ethylene-binding domain of ETRs seems to be essential for their function. The mutations located in the transmembrane domain of *SlETR1* (*Sletr1*-*1*), *SlETR3* (*Nr*), and its vicinity (*Sletr1-2*) confer dominant ethylene insensitivity in etiolated seedlings and delayed fruit ripening (**Okabe et al., 2011; Lanahan et al., 1994; Wilkinson et al., 1995**). Similarly, the mutations in the GAF domain of *SlETR5* (*Sletr5-1*) and its vicinity in *SlETR4* (*Sletr4-1*) influence the fruit ripening, with *Sletr5-1* fruits showing faster ripening (**Mubarok et al., 2019**). Distinct from these ETRs, *SlETR7* knockout and overexpression lines do not show any perceptible change in the fruit ripening process, though it affected the plant’s vegetative growth (**Chen et al., 2020).**

The cross-regulation of different ETRs has been observed, as in *Nr* fruits, mRNA and protein levels of other ETRs are affected relative to WT (**Chen et al., 2019; Hackett et al., 2000**). Among ETRs, SlETR3 and SlETR4 are the only receptors whose mRNA and protein abundance profiles follow the climacteric ripening process; both increase significantly at the onset of ripening (**Mata et al., 2018; Chen et al., 2019; Shinozaki et al., 2018**). Similar to SlETR3, SlETR4 is also critical for ripening regulation, and it is repeatedly phosphorylated at different levels depending on the ripening stage and ethylene action (**Kamiyoshihara et al., 2012**).

Unlike CTRs, which are like ETRs and are present as a multi-gene family, the tomato genome possesses only a single copy of *SlEIN2* (**Liu et al., 2015**). Consequently, the loss-of-function mutants of SlEIN2 result in yellow-fruited tomato plants and lack ethylene sensitivity (**Gao et al., 2016**; **Huang et al., 2022**). On the other hand, SlEILs being a multi- gene family, its mutant shows slower ripening, only as double or triple mutants (**Huang et al. 2022**).

In this study, we assessed the role of ethylene signaling in tomato seedlings and fruit ripening by analysing a loss-of-function ethylene-insensitive mutant isolated as an acetylene- resistant mutant. The mutation was localized in the GAF domain of the *ETR4* gene of tomato. The loss of function in ETR resulted in ethylene-insensitive seedlings and slower progression of fruit ripening. The retention of high lycopene and β-carotene levels post-red-ripe stage indicates that manipulation of ethylene signaling can be a potential target for improving the nutraceutical potential of tomato.

## Results

### Characterization of acetylene-resistant 1 (*atr-1)* mutant

The *atr-1* mutant was isolated by monitoring the absence of seedling triple response in the presence of acetylene (**Figure 1A**). It is a monogenic recessive mutant showing a Mendelian segregation ratio close to 3:1 (**Figure S1**). The dark-grown seedlings of the mutant showed reduced emission of ethylene than WT (**Figure S2A**). Consistent with acetylene mimicking ethylene, dark-grown *atr-1* seedlings did not show the triple response in the presence of ethylene (**Figure 1B**). Ethylene and 1-aminocyclopropane-1-carboxylic acid (ACC) treatment shortened the primary root and hypocotyl of WT seedlings almost to half-length of *atr-1* seedlings (**Figure 1C-D, Figure S2B**). The reduced sensitivity to the above treatments indicated that *atr-1* is an ethylene-resistant mutant. We next examined whether *atr-1* is also resistant to kinetin, exposure to an excess amount of which elicits overproduction of ethylene (**Cary et al., 1995**). Similar to ACC, *atr-1* seedlings at all dosages of kinetin had consistently longer hypocotyls than WT. Unlike hypocotyls, the exposure to kinetin drastically suppressed root elongation in both WT and *atr-1* seedlings (**Figure 1E, Figure S2C**).

**Figure 1.**
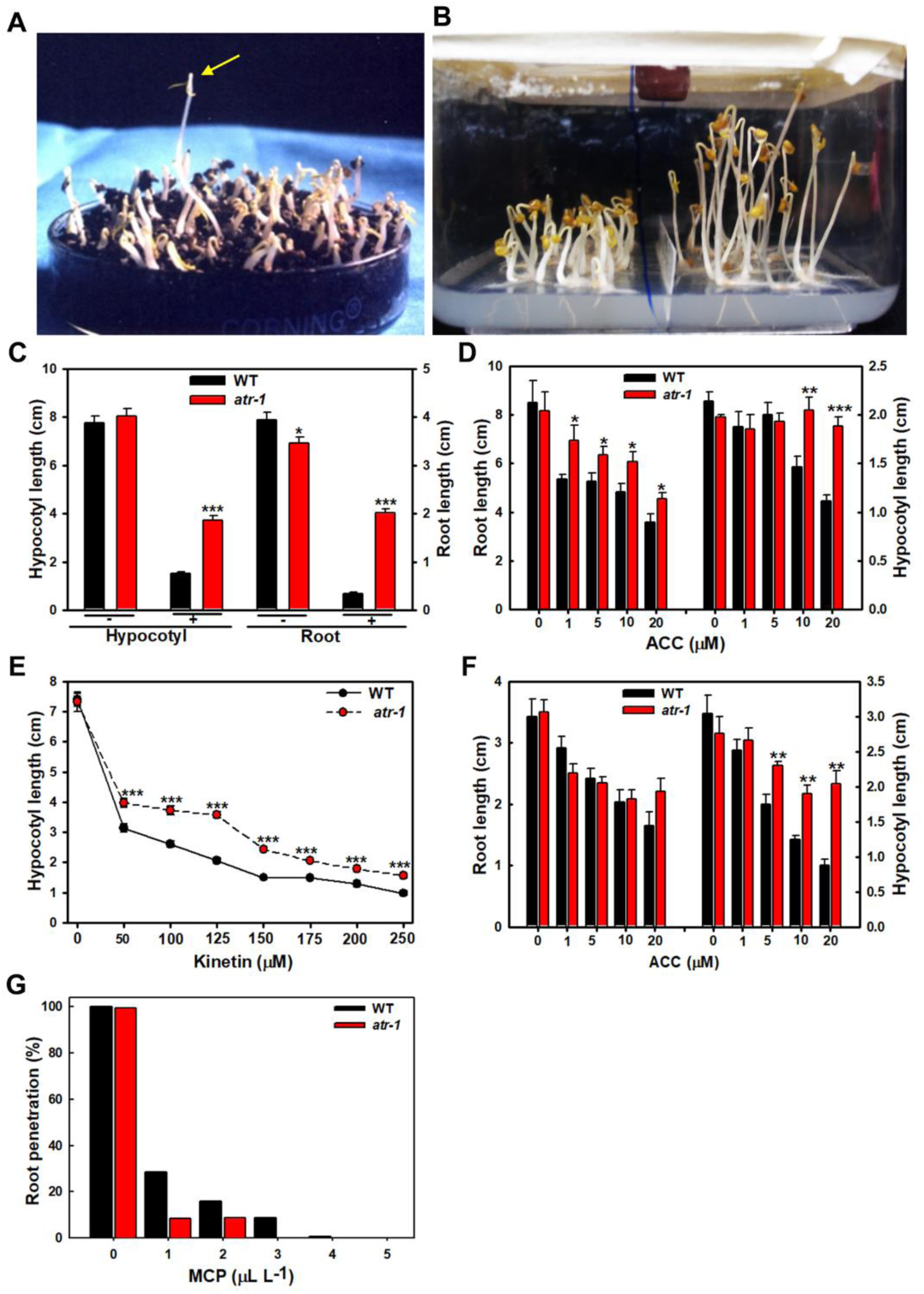
Characterization of *atr-1* mutant. (**A**) Isolation of *atr-1* mutant. The yellow arrow shows an acetylene-insensitive seedling, while other seedlings are dwarf and exhibit triple response. (**B**) Phenotype of WT and *atr-1* seedlings (7 days old) grown in the presence of ethylene (2 mL L^-1^). (**C**) Hypocotyl and root length of 5-day-old WT and *atr-1* seedlings grown without (**-**) or with (**+**) ethylene (2 mL L^-1^) Error bars show the standard error (±SE) for 25 seedlings, and each experiment was repeated more than three times. (**D**) Effect of ACC treatment on hypocotyl and root elongation of 5-day-old WT and *atr-1* seedlings. Error bars show the standard error for seven to ten seedlings, and each experiment was repeated more than three times. (**E**) Effect of varying concentrations of kinetin on the growth of 7-day-old etiolated seedlings of WT and *atr-1.* Error bars show the standard error for 40-45 seedlings, and each experiment was repeated more than three times. (**F**) Effect of ACC on hypocotyl and root elongation of 7-day-old light-grown seedlings. Error bars show the standard error for 7-10 seedlings, and each experiment was repeated three to four times. **(G)** Percent root penetration of 7-day-old *atr-1* and WT seedlings grown in different concentrations of 1-MCP. Asterisks indicate statistically significant differences between WT and *atr-1* seedlings. (Student’s t- test, * < 0.05, **< 0.01, ***< 0.001).

### *atr-1* shows reduced sensitivity to ethylene

In Arabidopsis, light-grown seedlings of ethylene-insensitive mutants show no or little increase in the hypocotyl length on exposure to ethylene (**Smalle et al., 1997**). The ACC inhibited hypocotyl elongation of light-grown WT seedlings but had little effect on *atr-1* hypocotyls. Conversely, ACC-mediated inhibition of root elongation was more severe in WT than *atr-1* ((**Figure 1F**, **Figure S3A**). We then examined whether the response to 1- methylcyclopropene (1-MCP), which tightly binds to ethylene receptors, such as the absence of root penetration in Soilrite (**Santisree et al., 2011**), is also altered in *atr-1*. 1-MCP elicited the loss of root penetration, albeit the *atr-1* was more sensitive. At 1.0 µL L^-1^ 1-MCP, only 10% of *atr-1* roots penetrated Soilrite compared to 30% for WT (**Figure 1G, Figure S3B**). One effect of ethylene is stimulation of senescence (**Iqbal et al., 2017**). Consistent being ethylene-resistant, multiple responses in *atr-1,* such as the dark-induced senescence of seedlings, ethylene-induced senescence of detached leaves, ethylene-triggered cotyledon, and floral abscission, were subdued compared to the WT (**Figure S4A-D**).

### *atr-1* seedlings show resistance to ABA

Since Arabidopsis ethylene-insensitive mutants display resistance to ABA (**Beaudoin et al. 2000)**, we examined the effect of ABA on the onset of seed germination, marked by the emergence of radicle, in *atr-1* and WT. In *atr-1,* the radicle emergence is delayed by a day (WT- 2^nd^ day; *atr-1* 3^rd^ day from sowing); however, by the 6^th^ day, WT and *atr-1* seeds are fully germinated. The effect of ABA manifested in a biphasic pattern, with phase I marking germination onset and phase II marking attainment of maximal germination. In a dose-dependent fashion, ABA severely affected both phase I and II in WT than *atr-1* (**Figure 2A**). The *atr-1* seedlings also showed lesser inhibition of hypocotyl and root growth than the WT with varying concentrations of ABA (**Figure 2B**, **Figure 5SA**). We then examined the effect of glucose that reportedly inhibits seed germination and seedling growth by elevating the ABA biosynthesis or ABA signaling (**Cheng et al., 2002; Finkelstein et al., 2002**). Similar to ABA, glucose also affected seed germination in a biphasic pattern. Both germination onset and attainment of maximal germination were more severely affected in WT than *atr-1* (**Figure 2C**). Since *atr-1* is an ethylene-resistant mutant, we compared its glucose resistance with the tomato *Never-ripe* (*Nr*) mutant, which has a truncated *ETR3* gene (**Figure 2D**, **Figure 5SB**). Contrasting to *atr-1,* which was resistant, the root growth of *Nr* was hypersensitive to increasing concentrations of glucose, showing severe inhibition of the elongation. Since the glucose-mediated inhibition of root elongation in *atr-1*, WT, and *Nr* is reverted by supplementing Norflurazon, an inhibitor of ABA biosynthesis (**Dejonghe et al., 2018**), the glucose induction of ABA seems to be a causal factor for the observed phenotype (**Figure 5SC**).

**Figure 2.**
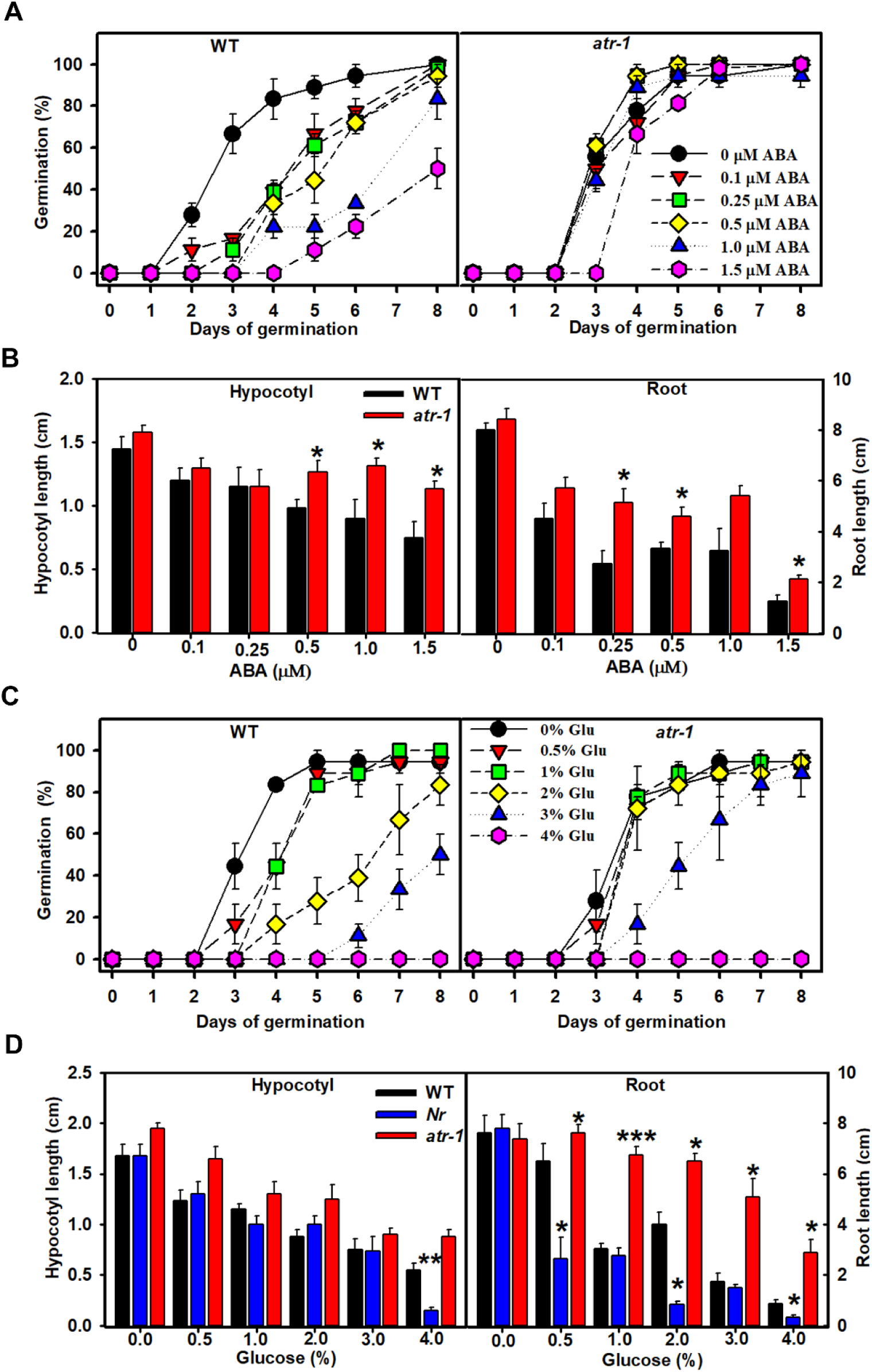
Effect of ABA and glucose on germination and seedlings growth of *atr-1* mutant. **(A)** Effect of different concentrations of ABA as indicated in the figure on germination of WT (left) and *atr-1* (right) mutant seeds. Error bars show the standard error (±SE) for eight seeds, and each experiment was repeated three times. (**B**) Hypocotyl (left) and root (right) length of 8-day-old light-grown *atr-1* and WT seedlings. Error bars show the ±SE for six seedlings, and each experiment was repeated three times. Asterisks indicate statistically significant differences between WT and *atr-1* seedlings. (Student’s t-test Student’s t-test, * < 0.05). **(C**) Effect of different glucose concentrations as indicated in the figure on WT (left) and *atr-1* (right) mutant seeds germination. Error bars show the standard error (±SE) for nine to 10 seeds, and each experiment was repeated three times. (**D**) Effect of glucose on hypocotyl and root growth 8-day-old WT, *Never-ripe (Nr),* and *atr-1* seedlings. Error bars show the ±SE for four to six seedlings, and each experiment was repeated three times (Student’s t-test, * < 0.05, **< 0.01, ***< 0.001).

### Exogenous ethylene alters the metabolite profile of *atr-1* seedlings

The air-grown etiolated seedlings of *atr-1* were phenotypically indistinguishable from the WT; however, on exposure to ethylene, growth inhibition in *atr-1* seedlings is milder than in the WT. The principal component analysis of the metabolome of air-grown *atr-1* and WT seedlings showed only mild separation between these two on PC1 (**Figure 3A**). The exposure to ethylene led to a shift in the PC of both WT and *atr-1* seedlings, the shift being milder for *atr-1* encompassing both PC1 and PC2, while for WT, it was mainly in PC1. Consistent with PCA, though ∼96 metabolites were detected, only a few metabolites (32) showed significant change in *atr-1* from WT treated with or without ethylene, and the affected metabolites were mainly downregulated (**Figure 3B**). The downregulated metabolites in *atr-1* mainly consisted of amino acids, and only a few overlapped with ethylene treatment. The ethylene treatment also led to massive downregulation of stress-related metabolites in *atr-1,* such as trehalose and dopamine. In air-grown *atr-1* seedlings, phytohormones IAA, ABA, SA, JA, and MeJA levels were higher than WT. The exposure to ethylene stimulated JA and ABA levels in *atr*-1 (**Figure 3C**).

**Figure 3.**
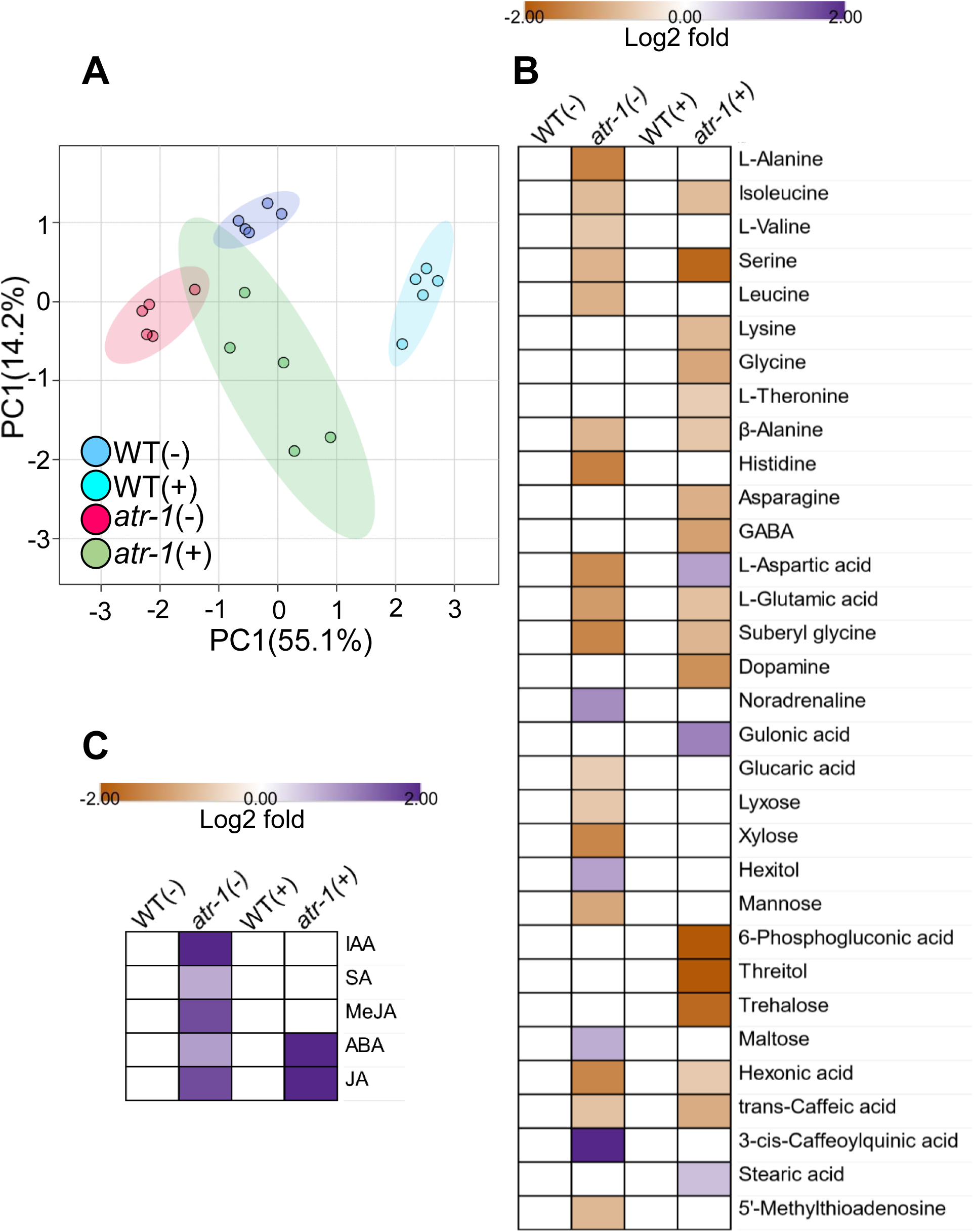
Metabolite and hormonal profiles of 5-day-old WT and *atr-1* seedlings grown with (+) and without ethylene (-). (**A**) Principle component analysis (PCA) of metabolites. The PCA was constructed using MetaboAnalyst 5.0. The variance of the PC1 and PC2 are given within parentheses. (**B-C**) The relative level of metabolites **(B**) and hormones (**C**) in *atr-1* seedlings. The relative changes in the metabolite (**B**) and hormone (**C**) levels were determined by calculating the log2 fold change. Only significantly changed metabolites (n ≥ 5) and hormones (n ≥ 3-5 [200 seedlings]) with reference to WT are depicted (Log2 fold ≥ ± 0.584, p-value ≤ 0.05). See **Dataset S1 and Dataset S2** for individual metabolite and hormone levels and significance, respectively. Abbreviation: ABA- Abscisic acid; IAA- Indole acetic acid; JA- Jasmonic acid; MeJA- Methyl Jasmonate; SA- Salicylic acid.

### The *atr-1* affected adult plant growth and fruit set

The *atr-1* mutant also influenced the vegetative and reproductive phases of the adult plant. In light-grown seedlings, primary leaves emerged earlier than WT. The leaves of the *atr-1* plant were bigger than WT leaves and had higher chlorophyll levels. One-month-old mutant plants were taller than WT. Consistent with this, the internodes in *atr-1* were longer than the WT (**Figure S6A)**. The *atr-1* had almost double the number of flowers per inflorescence truss compared to WT. Most flowers in *atr-1* set fruits, resulting in an increased fruit set. The average number of fruits per truss in *atr-1* was (6±1), whereas each truss of WT had (3±1) fruits (**Figure 4A-F**). The fruit size of WT and *atr-1* was nearly the same; however, *atr-1* fruits had fewer seeds (WT∼98; *atr-1*∼50).

**Figure 4.**
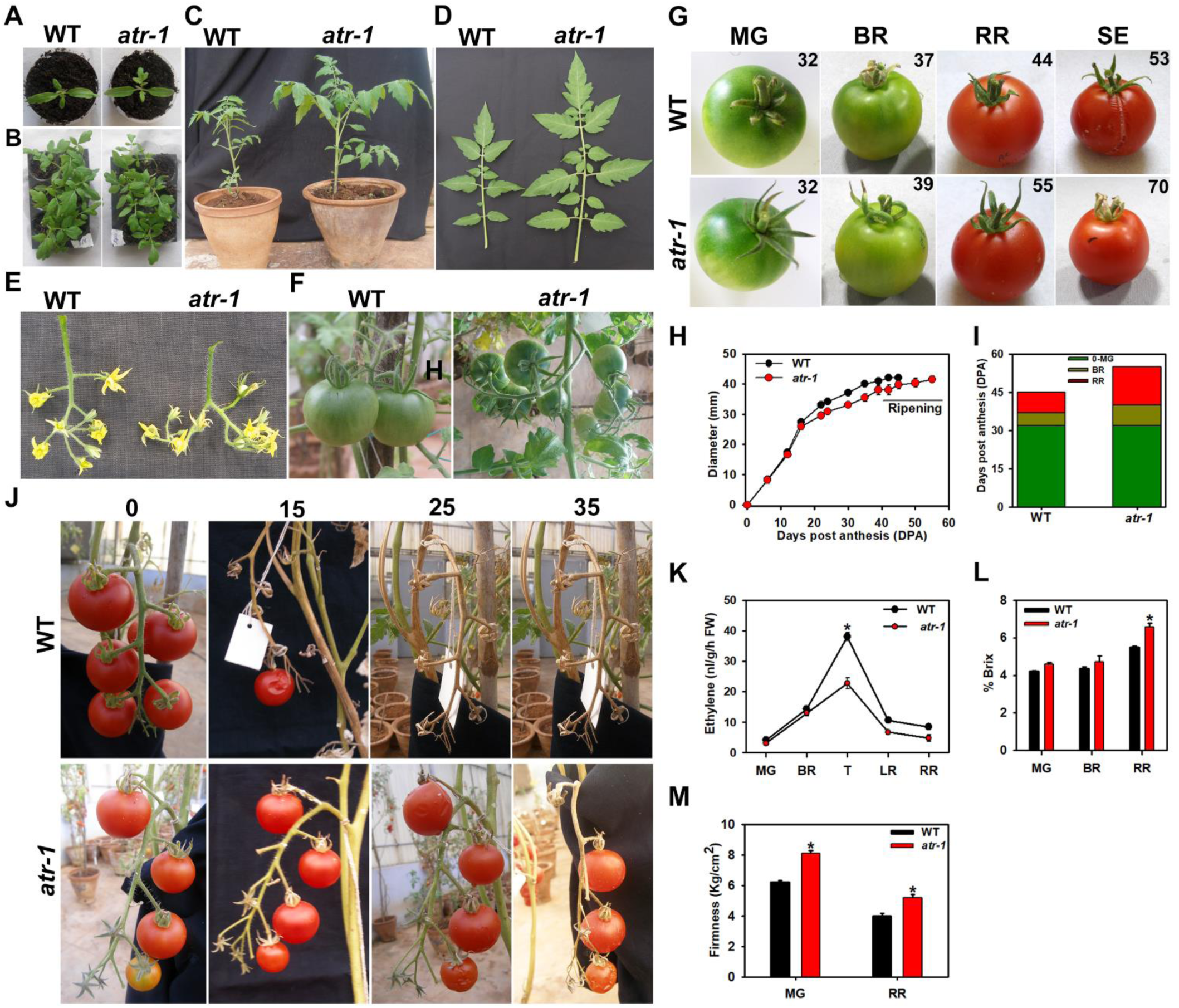
The *atr-1* mutant shows an altered phenotype throughout the life cycle. (**A, B**) Picture of 7- (**A**), 15- (**B**) days-old WT and *atr-1* seedlings grown in a growth chamber under white light. (**C**) Morphology of 30-days-old WT and *atr-1* plants grown in greenhouse. (**D**) *atr-1* mutant shows broad leaflets in comparison to WT. Leaf was harvested from the 7^th^ node of one-month-old plants. (**E, F**) the increased number of flowers (**E**) and fruits (**F)** per inflorescence of *atr-1* compared to WT.(**G**). Fruits of WT and *atr-1* were harvested at different stages of ripening. The post-anthesis age of the fruit in days is indicated in the top right corner of the photos. **MG**-mature green, **BR-**breaker, **RR**- red ripe, **SE-**senescence (**H**) Time course of increase in fruit diameter from anthesis to the red ripe stage. (**I**) Chronological development of WT and *atr-1* fruits from anthesis to fruit senescence. **LR**- light red. Each bar stack in the ripening graph corresponds to an average of seven fruits from the first truss of wild type and mutant. (**J**) RR fruits were examined for on-vine senescence. The post-RR age of fruits in days is indicated on top of the photos. (number of plants, n ≥ 5). (**K**) Ethylene emission from WT and *atr-1* fruits at different stages of ripening. (**L**) % Brix level in WT and *atr-1* fruits at MG, BR, and RR stages. (**M**) Firmness of *atr-1* and WT fruits at MG and RR stage. Asterisks show statistically significant differences between WT and *atr-1.* (Student’s t-test, *P < 0.05, number of fruits, n ≥ 5 *±* SE) **MG**-mature green, **BR**-breaker, **T**-turning, **LR**-light red, **RR**-red ripe.

### Ripening duration is extended in *atr-1*

Post-anthesis, the growth of *atr-1* fruits was nearly parallel with the WT. Though *atr- 1* and WT attained the mature-green (MG) stage at the same time (32 DPA), the duration to reach the breaker stage (BR) was slightly longer in *atr-1*, whereas the duration to reach the red-ripe (RR) stage was much longer in *atr-1* than in WT. While WT fruits attained the RR stage in 6-7 days, *atr*-1 fruits required 13-15 days to reach the RR stage. The *atr-1* mutation also influenced the on-vine longevity of the fruits. Post-RR stage, the *atr-1* fruits showed senescence symptoms after 25-30 days, while WT fruits dropped from the vine within 15 days (**Figure 4G-J**). The red-ripe *atr-1* fruits were firmer than WT and had a higher %Brix. Similar to seedlings, ethylene emission from the *atr-1* fruits was lower than the WT, particularly at the turning (TUR) stage (**Figure 4K-M**). The post-harvest shelf life of a*tr-1* fruits detached at the RR stage was similar to WT, with symptoms of senescence manifested as skin wrinkling appearing at the same time. However, the detached *atr-1* RR fruits were more resistant to high temperatures (45°) than the WT (**Figure S6B-D**).

### **β**-carotene level is sustained in *atr-1* fruit post-RR Stage

The carotenoid profile of the *atr-1* fruit was also different from the WT (**Figure 5, Figure S7**). Particularly, post-RR stage, while lycopene and β-carotene levels declined in WT fruits, both were retained at higher levels in post-RR fruits of *atr-1* for nearly 15 days. Notably, the β-carotene level in *atr-1* was higher than WT at the RR stage, which was sustained post-RR stage for almost 15 days. The retention of higher β-carotene levels in the *atr-1* post-RR stage was associated with its precursor γ-carotene, which was also maintained at a higher level. Another feature was the detection of phytofluene in *atr-1* fruit post-RR, suggesting that carotenoid synthesis continues in *atr-1* even after the attainment of full ripening. Contrasting to phytofluene, phytoene in *atr-1* was detected only at the RR+7-day stage. In WT, the phytoene was first detected at RR, and its levels declined post-RR stage. Though the lutein level gradually declined in *atr-1* and WT, its levels were detectable till the RR+15D stage. In contrast, neoxanthin and violaxanthin declined to a very low level at the RR stage.

**Figure 5.**
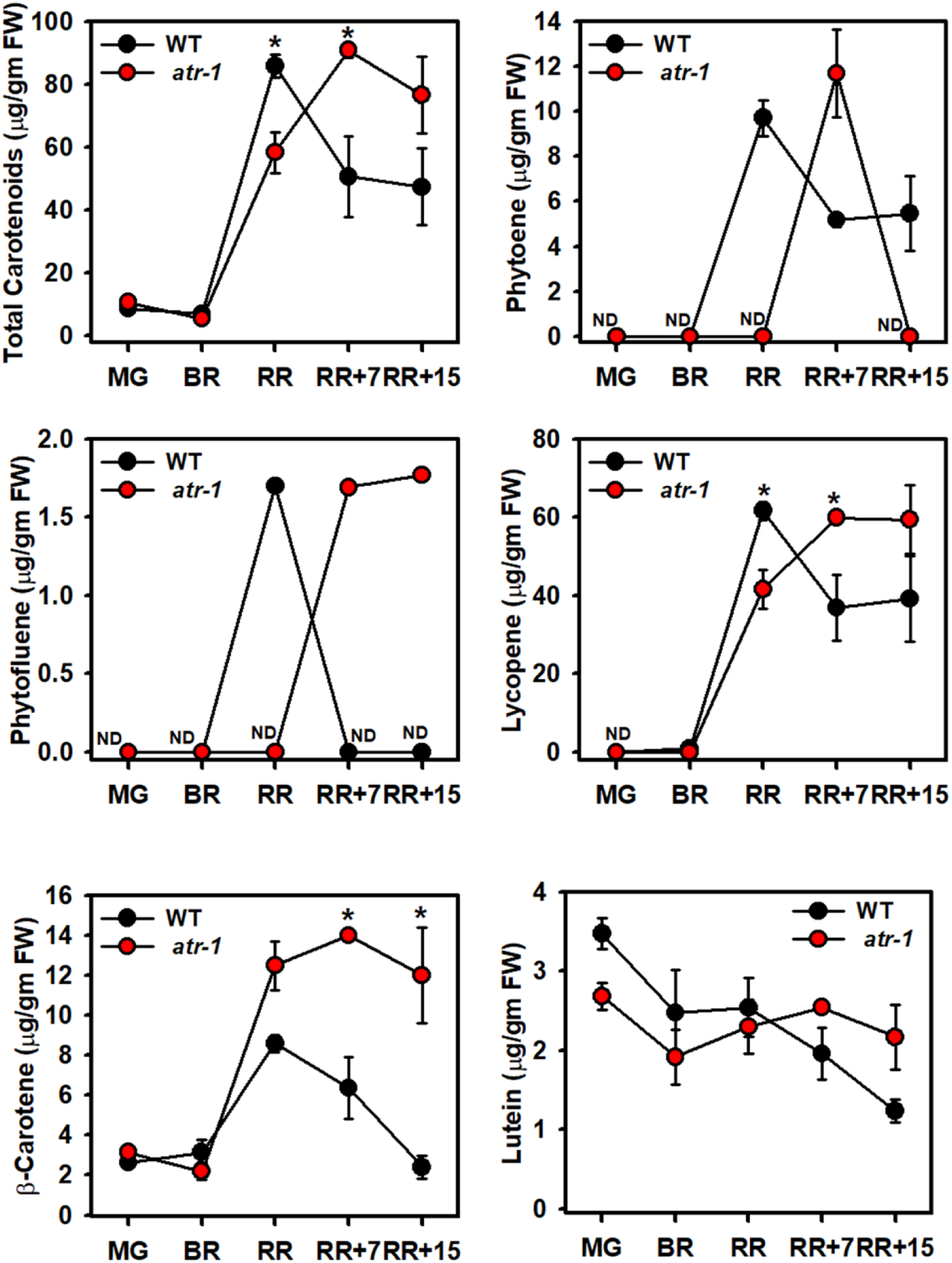
The time course of increased in different carotenoid levels in *atr-1* and WT (AC) fruits. The carotenoid data are expressed as mean ±SE (n ≥ 3). *P ≤ 0.05**. See Dataset S3** for individual carotenoid levels and significance. ND- not detected

### The metabolome of *atr-1* fruit differed in a stage-specific pattern

Unlike seedlings, the ethylene emission from *atr-1* fruits was lower than WT only at the TUR stage of the ripening. Though the onset of ripening was delayed in *atr-*1, it did not display a significant shift in levels of phytohormones during fruit ripening. While MeJa at MG, IBA and ABA at BR, and JA levels were lower at RR stages respectively, the RR+15D fruits had high MeJA and SA levels (**Figure 6A**). No phytohormone was significantly changed at the RR+7D stage than WT. Seemingly reduced ethylene sensitivity of the *atr-1* mutant did not influence the levels of other phytohormones.

**Figure 6.**
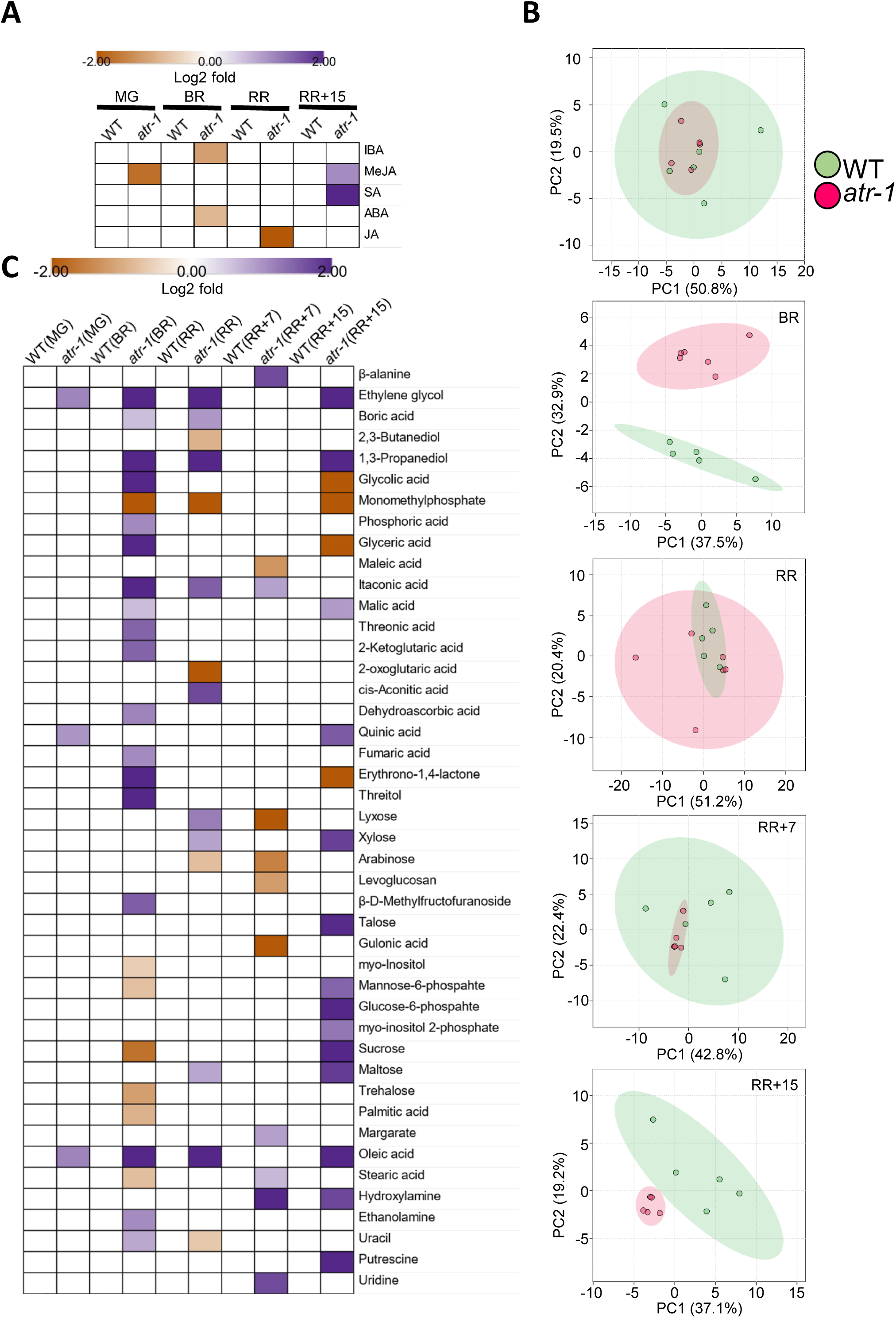
Metabolite and hormone profiles WT and *atr-1* fruits at different ripening stages. (**A**) Relative levels of hormones during fruit ripening. Only significantly changed hormones with reference to Ailsa Craig (WT) are depicted (Log2 fold ≥ ± 0.584, p-value ≤ 0.05, n ≥ 3). At RR+7 days, since no hormone was significantly changed, this stage is not depicted in the heatmap. See **Dataset S4** for individual hormone levels and significance. *Abbreviation*: MeJA- Methyl Jasmonate, IBA- Indolebutyric acid, ABA- Abscisic acid, JA- Jasmonic acid, SA- Salicylic acid. (**B**) Principle component analysis (PCA) of metabolites. The PCA was constructed using MetaboAnalyst 5.0. The variance of the PC1 and PC2 are given within parentheses. (**C**) The relative level of metabolites in *atr-1* fruits. The relative changes in the metabolite levels were determined by calculating the log2 fold change. Only significantly changed metabolites, with reference to WT at the respective stage, are depicted (Log2 fold ≥ ± 0.584, p-value ≤ 0.05, n ≥ 5). See **Dataset S5** for metabolite levels and significance.

The metabolite profiles of the *atr-1* fruit differed from WT in a stage-specific fashion during fruit ripening. The principal component analysis (PCA) of the metabolite profile did not show a major difference at MG. However, at BR, the metabolome of *atr*-1 showed a distinct shift away from WT, particularly for PC2. At the RR stage and RR+7 days stage, the PCA of *atr-1* and WT largely overlapped, with WT and *atr-1* showing a more compact distribution at RR and RR+7D, respectively. A distinct separation of PCA was again observed at the RR+15D stage, wherein *atr-1* differed from WT, particularly for PC1 (**Figure 6B**).

Comparison of significantly changed metabolites with respect to WT at each ripening stage showed a dynamic shift in relative expression levels of metabolites. Consistent with an overlap in PCA at the MG stage, WT and *atr-1* fruit did not display a major difference except for levels of three metabolites. Among the above three metabolites upregulated at the MG stage, two of them, ethylene glycol and oleic acid, were also at higher levels at the BR, RR, and RR+15-day stages (**Figure 6C**). Contrasting to MG and consistent with PCA at the BR stage, a major shift in metabolite levels was observed in *atr-1* at the BR stage. Out of 82 metabolites detected in tomato fruits, 25 metabolites showed significant change at BR. Notably, the metabolites related to and derived from the TCA cycle, such as itaconic acid, malic acid, 2*-*ketoglutaric acid, and fumaric acid, were upregulated. The stress-related compound trehalose was downregulated at BR.

Compared to BR, relatively few metabolites (14) were modulated in *atr-1* at RR Stage. Here, too, metabolites related to the TCA cycle, such as itaconic acid and cis-aconitic acid, were upregulated. Also, the sugars such as xylose and maltose were upregulated. Like RR, at the RR+7D stage, only a few metabolites were affected in *atr-1*. However, the stress-related compound hydroxylamine was upregulated at RR+7D and RR+15D stages. Similar to BR, as reflected in PCA, more prominent metabolite changes were observed at the RR+15D stage. At the RR+15D stage, prominently sugars such as xylose, sucrose, maltose, talose, mannose- 6-phosphate, and glucose-6-phosphate were upregulated.

### Candidate genes

As the growth of *atr-1* mutant seedlings is resistant to ethylene, it can be surmised that it is mutated in either in ETRs or a downstream signal transduction pathway component like EIN2 or EILs (**Okabe et al., 2011; Huang et al. 2022**). The recessive nature of the *atr-1* loci indicated that it was a weaker allele. The semi-quantitative PCR of the *ETR* genes in seedlings and fruit ripening showed expression levels nearly similar to the WT (**Figure S8**). This indicated that mutation in *atr-1* loci did not alter the expression of the *ETR* gene family in tomato. Among the downstream genes, the *EIN2* gene regulating ethylene signaling also serves as a central point for regulating tomato ripening (**Gao et al., 2016; Huang et al., 2022**). The sequence analysis and digestion with a mismatch-specific endonuclease CEL-I revealed no mutation in the entire ORF and the promoter of *EIN2* of the *atr-1* mutant (**Figure S9**). Likewise, no mutation was found in tomato *EIL1*, *EIL2,* and *EIL3* genes of *atr-1* (**Figure S10**).

To decipher the candidate gene in *atr*-1, we sequenced and compared its whole genome with the parental line Ailsa Craig. Based on the SIFT score, which predicts whether or not a mutation is deleterious, 371 unique genes were predicted to bear the deleterious mutation (**Dataset S6**). Most of these genes belonged to the multi-gene families and had a structural function; therefore, they may not have a wide-ranging effect, such as the ethylene resistance of *atr-1*. Among ethylene receptors and downstream signaling components, a deleterious mutation P279L was detected in the *ETR4* (Solyc06g053710) gene. Tomato ETR4 is a member of the ethylene receptor subfamily II and homologous to AtEIN4. The ethylene receptor consists of a membrane-localized N-terminal ethylene-binding domain and a C-terminal cytosolic receptor domain comprising a GAF domain followed by an HK and a receiver domain. The P279L mutation in the *ETR4* gene lies in the GAF domain (182-342aa) (**Figure 7A, Figure S11**). The proline in the GAF domain of ETR4 is a highly conserved amino acid in a wide range of plant species (**Figure S12**).

**Figure 7.**
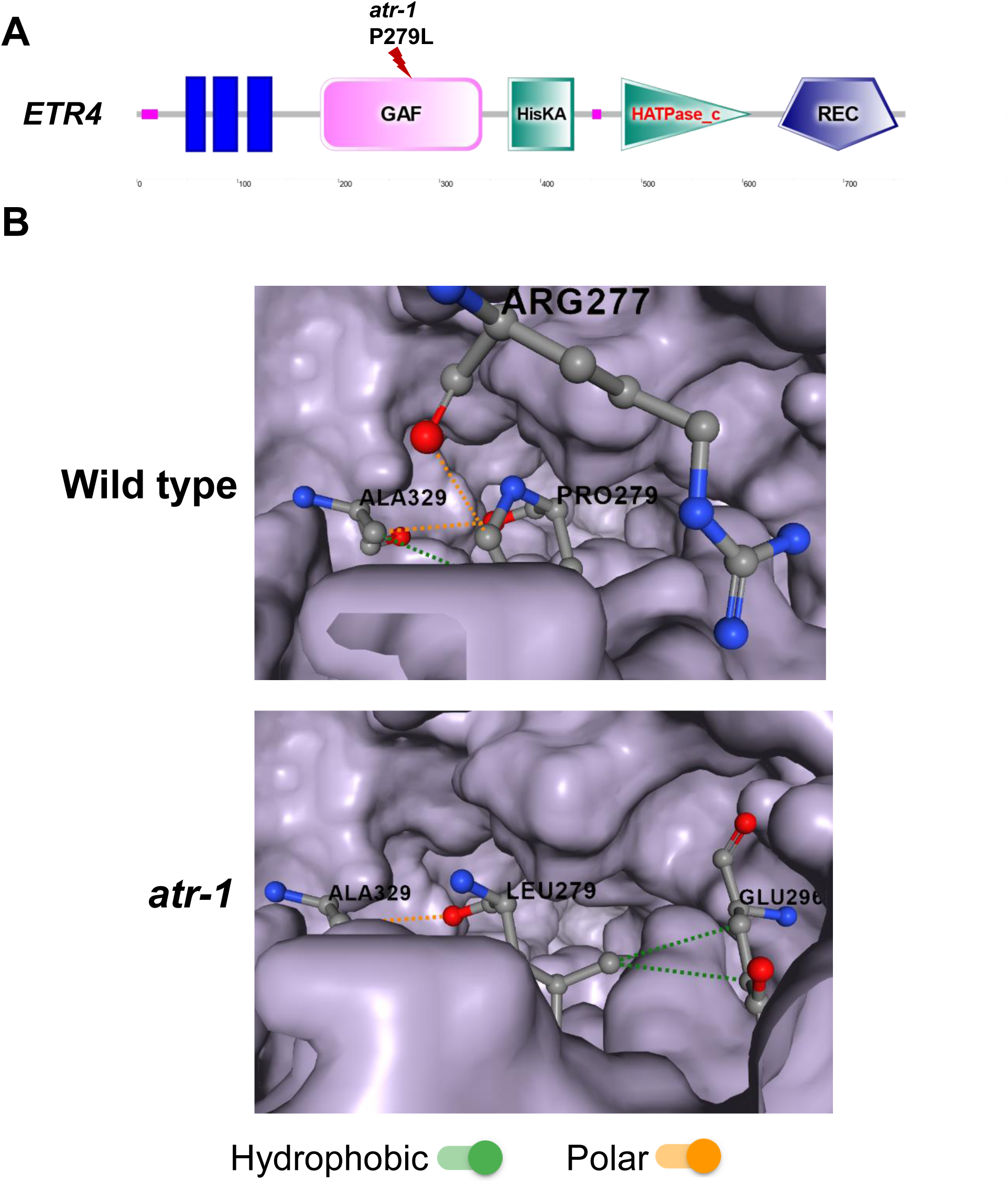
Location of mutation in the GAF domain of *atr-1* and its influence on protein structure**. (A)** The SMART (http://smart.embl-heidelberg.de/) image of the ETR4 protein showing the location of the mutation in the GAF domain in *atr-1*. The transmembrane domain, GAF domain, histidine kinase A domain, histidine kinase-like ATPase, and receiver domains are shown in the cartoon (**Dataset S6**). (**B**) The modeling of ETR4 by DDmut (https://biosig.lab.uq.edu.au/ddmut). In WT, proline 279 has a hydrophobic and polar interaction with alanine 329 and a hydrophobic interaction with arginine 327. In *atr-1,* the interactions have changed. Leucine 279 has a polar interaction with alanine 329 and two hydrophobic interactions with glutamine 296. The interaction with Arginine 277 is absent in the *atr-1* mutant. This shift in interaction is predicted to have a stabilizing effect [Gibbs free energy ΔΔG 0.25 kcal/mol) on *ETR4* protein encoded by *atr-1*.

The studies on protein-protein interactions have suggested that dimer formations in the ETR family are substantively mediated by the GAF domain (**Grefen et al., 2008; Berleth et al., 2019**); therefore, P279L mutation in *atr-1* may affect ETR4 receptor function (**Figure 7A**). The mutant ETR4 protein modelling indicated that the mutation slightly increases Gibbs Free Energy (ΔΔG = 0.25 kcal/mol). Though the mutation promotes protein stability, the mutated amino acid leucine 279 has altered interaction with neighboring amino acids (**Figure 7B**). The above shift in amino acid interactions in mutated ETR4 may influence the GAF domain interaction with other proteins.

## Discussion

### The *atr-1* is an ethylene-resistant mutant

The phenotype of *atr-1* is reminiscent of Arabidopsis *Atetr* and *Atein* mutants, as its dark-grown seedlings lacked triple response on exposure to ethylene **(Guzman and Ecker, 1990**). However, the *atr-1* seedlings were only partially resistant to ethylene. The ethylene exposure elicited inhibition of hypocotyl and root growth in *atr-1*, however, the inhibition was notably milder than the WT. Consistent with the above, the resistance to growth inhibition in hypocotyl and root of *atr-1* seedlings was manifested only at a higher dosage of ACC (precursor of ethylene). Like dark-grown seedlings, the light-grown seedlings of *atr-1* were also resistant to growth inhibition, albeit the resistance of root was manifested at higher ACC dosage. The resistance of *atr-1* seedlings was not limited to exogenous ethylene; its hypocotyl growth was also resistant to kinetin, which suppresses seedling growth by stimulating endogenous ethylene biosynthesis (**Hansen et al., 2009**). Similar to *atr-1*, tomato *Sletr1-2* and *Sletr4-1*dark-grown seedlings also showed lesser growth inhibition than WT on exposure to ethylene (**Okabe et al., 2011; Mubarok et al., 2019**). Contrariwise, dark-grown seedlings of the *Sletr1-1* and *Slein2-1* mutants were fully resistant to ethylene (**Okabe et al., 2011; Huang et al., 2022**). Since mutations in *Sletr1-1* and *Sletr1-2* are located in different regions of the *Sletr1* gene, this may be the reason for the differential sensitivity of these two alleles to ethylene (**Okabe et al., 2011**), while total loss of ethylene-sensitivity in *Slein2-1* is related to EIN2 being a single copy in tomato genome (**Huang et al. 2022**).

Since *atr-1* is similar to tomato *Sletr1-2* and *Sletr4-1* mutants that are less sensitive to ethylene than WT, it is plausible that the ethylene sensing response may have been attenuated in *atr-1*. Tomato *Sletr3* (*Nr*) mutant shows loss of root penetration in Soilrite, which is exacerbated by 1-MCP, an inhibitor of ethylene receptors (**Santisree et al., 2011**). Since 1- MCP-mediated inhibition of root penetration in Soilrite was more pronounced in *atr-1* than WT, *atr-1* seems to be compromised in ethylene perception. The likelihood of reduced ethylene perception by *atr-1* also manifests in detached leaf senescence, where *atr-1* leaves incubated with or without ethylene show delayed senescence than WT, like *slein2-1* detached leaves (**Huang et al., 2022**). The resistance to cotyledon and flower abscission in *atr-1* seedlings and inflorescences exposed to ethylene conform to reduced ethylene perception by *atr-1*. The reduced ethylene emission from *atr-1* seedlings might be related to reduced ethylene perception, as vegetative tissue shows the autocatalytic induction of ethylene emission (**Riov and Yang, 1982**), which may be compromised in *atr-1*.

### Interaction with ABA is affected in *atr-1*

The hormonal modulation of plant development involves close interaction among different hormones. Among these, during seed germination, ethylene, GA, and ABA play a prominent role (**Linkies and Leubner-Metzger, 2012).** The delay in the onset of *atr-1* seed germination likely emanates from reduced ethylene perception, as reduced ethylene emission from tomato *acs2-2* mutant also delays the seed germination (**Sharma et al., 2021**). Contrary to the WT, where exogenous ABA suppresses germination, *atr-1* seeds were resistant to ABA-mediated inhibition of germination. Similar to ABA, the exogenous glucose also suppresses seed germination, likely by interfering with ABA catabolism (**Price et al., 2003**). Since *atr-1* is also resistant to glucose-mediated germination inhibition, it indicates an overlap between ABA and glucose responses. Seemingly, glucose stimulates ABA synthesis as the inclusion of Norflurazon, an ABA biosynthesis inhibitor (**Dejonghe et al., 2018**), reverses the inhibitory effect of glucose.

The *atr-1* resistance to ABA is opposite to Arabidopsis ethylene-insensitive *Atein2-1* and *Atetr1-1* mutants, where exogenous ABA severely inhibited seed germination (**Beaudoin et al., 2000; Ghassemian et al., 2000**). Seemingly, the ethylene-insensitivity of *atr-1* renders it resistant to ABA for seed germination and seedling growth. It entails that the ethylene-insensitivity of *atr-1* is likely conferred by a signaling component other than one homologous to AtETR1 and AtEIN2. Current evidence indicates that AtETR1 and AtEIN2 regulate the ABA sensitivity of Arabidopsis seed germination independent of the canonical ethylene pathway by regulating ABA-responsive gene expression (**Guo et al., 2023; Bakshi et al., 2018**). It remains to be determined whether *atr-1* insensitivity to ABA is related to reduced expression of ABA-responsive genes or has an alternate mechanism.

### Ethylene insensitivity attenuates the metabolome profile of *atr-1* seedlings

Morphologically, *atr-1* seedlings grown in the air were indistinguishable from the WT. Little is known about the influence of ethylene on seedlings’ metabolome and phytohormones. The PCA of air-grown and ethylene-treated *atr-1* seedlings was distinct from WT. Most metabolites were downregulated in the air-grown *atr-1* seedlings, with amino acids constituting the most prominent class. The lower levels of metabolites in *atr-1* seedlings are akin to the ethylene underproducer tomato *acs2-2* mutant, where in leaves, most of the metabolites were downregulated (**Sharma et al., 2021**). Seemingly, the ethylene insensitivity of the *atr-1* seedling leads to altered metabolic homeostasis.

The exposure to ethylene shifted the metabolic profile of *atr-1* seedlings; most of the metabolites were downregulated. Similar to air-grown seedlings, the major affected group was amino acids. Mainly, amino acids linked with glycolytic pathway, such as lysine, glycine, and threonine, were lower. Lowering of amino acids may result from reduced biosynthesis or efficient incorporation in proteins. Since ethylene-treated *atr-1* seedlings were taller than WT, enhanced metabolic flux directed to the growth may be the reason for the lower metabolite levels. Consistent with this, the levels of stress-related metabolites such as trehalose, threitol, and dopamine (**Kumari et al., 2022**) were lower in ethylene-treated *atr-1* seedlings.

The ethylene-insensitivity of *atr-1* also affected the hormonal homeostasis manifested by increased levels of all phytohormones. Since the expression of the *NCED3* gene, a key enzyme in ABA biosynthesis, is upregulated in the *Atein2-1* mutant, it suggests that impairment of ethylene signaling enhances ABA biosynthesis (**Cheng *et al*., 2009**). Unlike IAA, MeJA, and SA, the ethylene treatment could not reverse ABA and JA upregulation. It remains to be deciphered how hormonal and metabolome changes are liked with ethylene- resistant growth of *atr-1* seedlings.

### Fruit ripening is slower in the *atr-1* mutant

Morphologically, *atr-1* plants were distinct from WT plants due to their faster growth and bigger leaves. Since ethylene-insensitive tomato *Nr* (*ETR3*) mutant plants are also taller than the WT (**Nascimento et al., 2020**), the faster growth of *atr-1* is likely related to its ethylene-insensitivity. The taller phenotype of *atr-1* and *Nr* plants seems to be specific to them as other ethylene-insensitive mutants of tomato viz. *Sletr1-1*, *Sletr1-2*, *Sletr4-1*, *Sletr5- 1,* and *slein2-1* show normal morphology like WT (**Okabe et al., 2011; Mubarok et al., 2015, 2019; Huang et al., 2022**). An increased number of flowers in the inflorescence could have aided in a higher fruit set of *atr-1* (**Soyk et al., 2017**), or it could be related to ethylene insensitivity as *Slein2-*1 also has a higher fruit set and reduced seed number in fruits (**Huang et al., 2022**).

The WT and *atr-1* fruits attained the MG stage at the same time. Seemingly, reduced ethylene sensitivity does not affect the duration needed to reach the MG stage in tomato. Consistent with this, the duration to reach the BR stage was not affected in other ethylene-insensitive *Sletr4-1* and *Sletr5-1* mutants of tomato (**Mubarok et al., 2019**). Conversely, unlike other ethylene-insensitive mutants (**Mubarok et al., 2019**), post-MG stage, the duration to reach the BR and RR stage in the *atr-1* was much longer. It is plausible that reduced ethylene emission from *atr-1* fruits and reduced ethylene sensitivity leads to slower ripening progression, as reported for combinations of *Sleils* mutants (**Huang et al. 2022**). Consistent with this, the ripening progression is slower in tomato *acs2-2* mutant, which, like *atr-1,* has reduced ethylene emission in ripening fruits (**Sharma et al., 2021**). Though the on-vine senescence and fruit drop were considerably delayed, the *atr-1* fruits harvested at RR, like *Sleils* mutants (**Huang et al., 2022**), did not display any increase in shelf life over WT. In contrast, tomato *Sletr1-2* and *Sletr4-1* mutants had improved post-harvest shelf life than WT (**Okabe et al., 2011; Mubarok et al., 2015, 2019).**

### *atr-1* fruits retain high lycopene and **β**-carotene levels post-RR stage

The onset of fruit-specific carotenogenesis is closely linked with ethylene biosynthesis and perception. The antisense suppression of *ACS2* (**Theologis et al., 1993**) or defects in ethylene perception in the *Nr* mutant (**Lanahan et al., 1994**) inhibit the accumulation of carotenoids in fruits. The slower accumulation of total carotenoids in *atr-1* fruit is consistent with its ethylene insensitivity. Among the carotenoids, *atr-1* accumulated significantly higher levels of β-carotene than WT. Moreover, the ethylene biosynthesis is also subdued in *atr-1* fruits, at least at the turning stage. Notably, the β-carotene was retained at high levels post-RR stage, while it declined in the WT. Commensurate with β-carotene, its precursor γ-carotene was also retained at a higher level. Since lycopene levels remain high in the *atr-1* post-RR stage, ethylene insensitivity of *atr-1* may be a causal factor sustaining high β-carotene derived from high lycopene levels.

The higher lycopene and β-carotene levels in *atr*-1 fruit are likely related to delayed senescence of on-vine attached fruits. Unlike WT, whose fruits drop from the vine ∼15 days from the post-RR stage, the *atr-1* fruits stay on the vine for nearly 30 days. The delayed senescence of *atr-1* fruits is likely related to the reduced sensitivity to ethylene. Since fruits of *acs2-2,* an ethylene underproducer mutant, also show delayed senescence (**Sharma et al., 2021**), it supports reduced ethylene insensitivity being the causal reason for the delayed fruit drop of *atr-1* fruits.

The metabolic profiles of *atr-1* fruits conform to this notion. The significant difference in metabolic profiles of WT and *atr-1* are seen at the BR and RR+15D stage. The BR stage is marked by an overall surge in metabolic activity with higher TCA cycle intermediates such as fumaric acid, malic acid, and 2-ketoglutaric acid. Conversely, increased levels of sugars are seen at the RR+15D stage, including sucrose and glucose-6-phosphate. In tomato DFD accession having a long shelf life, high β-carotene levels are associated with increased sucrose levels (**Osorio et al., 2020**).

### Mutation in the GAF domain reduces the ethylene sensitivity of *atr-1*

In tomato, mutations in ethylene receptor genes fall into two categories: dominant and recessive. The dominant mutations in *Sletr1-1*, *Sletr1-2*, and *Nr* (*SlETR3*) confer ethylene insensitivity in seedlings (**Okabe et al., 2011; Lanahan et al., 1994; Figure S13, Table S2**). These mutations are located in the ethylene-binding transmembrane domain. In Arabidopsis, single amino acid substitutions in the transmembrane ethylene-binding domain of any of five ethylene receptors result in dominant ethylene insensitivity (**Hall et al., 1999**). In *Curcurbita pepo, Cpetr1a* mutation in the transmembrane ethylene-binding domain is semi-dominant and displays ethylene insensitivity, whereas *Cpetr2b* mutation between the GAF and the histidine-kinase domain is also semi-dominant and ethylene-insensitive (**García et al., 2020**). Considering that the ethylene binding in ETRs is confined to the transmembrane domain, it is likely that the mutations in other regions of ETR, such as in *Cpetr2b,* interfere with the transduction of the ethylene signal.

Consistent with this, a mutation in the GAF domain of *atr-1* likely led to ethylene insensitivity by affecting the output of ethylene-mediated signaling. While the mutations in the ethylene-binding domain are dominant in nature (**Okabe et al., 2011; Lanahan et al., 1994**), the mutation in the GAF domain or its vicinity result in recessive mutants like *atr-1*, *Sletr4-1,* and *Sletr5-1* in tomato (**Mubarok et al., 2019, Figure S13, Table S2**). The GAF domain plays a role in heteromeric interactions between ethylene receptors (**Gao et al., 2008**). A direct role of the GAF domain in the regulation of tomato ripening is highlighted by using an archetypical inhibitory octapeptide, known as NOP-1 (LKRYKRRL), which plausibly hampers the receptor signal transduction to kinase and receiver domains (**Kessenbrock et al., 2017; Mili**ć **et al., 2018).** NOP-1 directly interacts with two ripening regulating ethylene receptors in tomato fruits, the SlETR3 (Nr) and SlETR4, and delays the ripening process.

It is plausible that in *atr4-1,* the mutation in the GAF domain influences the heteromeric interaction with other coexpressed ETRs. In ripening tomato fruits, SlETR4, though it belongs to subfamily II of ETRs, preferentially forms heteromers with subfamily I rather than with subfamily II members. SlETR4 interaction is not limited to ETRs; it also interacts with SlCTR1 and SlCTR3 directly or via ETRs; thus, the ethylene insensitivity of *atr-1* may ensue from its wide range of interactions (**Kamiyoshihara et al., 2022**).

Emerging evidence has indicated that SlETR4 regulates tomato ripening in concert with SlETR3. The relationship between these two is indicated by transcriptome analysis, where levels of *SlETR3* and *SlETR4* transcripts parallelly upregulate at the onset of ripening and are present at higher levels than the other *ETR* genes (**Figure S14**, **Shinozaki et al., 2018**). A direct interaction between the SlETR3 and SlETR4 is highlighted by reduced dephosphorylation of SlETR4 in *Nr* mutant on ethylene exposure. The SlETR3 seems to be the dominant partner, as in the transgenic SlETR3-suppressed line, the SlETR4 protein level is reduced, though its transcript level remains similar to the wild type (**Kamiyoshihara et al., 2022**).

The differences between the ripening behaviour of *atr-1* (P279L, GAF domain) and *Sletr4-1* (G154S, between the transmembrane and GAF domain, **Mubarok et al., 2019, Figure S13**) mutants may ensue from the mutations being located at different domains of SlETR4. The major difference between the two was that *atr-1* fruits had higher brix than WT and did not show any perceptible increase in the post-harvest shelf life over WT (2 days in *Sletr4-1*). In *Sletr5-1* (R278Q), the mutation in the GAF domain increases sensitivity to ethylene; fruit has a smaller size and shorter shelf life (**Mubarok et al., 2019**). Ostensibly, between different ETRs and regions of the GAF domain, the location of mutation affects the fruit phenotype differentially. It entails that mutation in the different regions of the GAF domain may help decipher its role in ethylene signaling in tomato fruit ripening.

A major target of tomato breeding programs is to prevent the deterioration of the nutrient content during the postharvest shelf life (**Osorio et al., 2020**). Unlike the *Nr* mutant, which interferes with ripening, leading to yellow fruits (**Lanahan et al. 1994**), the *atr-1* mutant shows the normal progression of the ripening. Notably, the retention of lycopene and β-carotene in *atr-1* fruits brings forth the continued role of ethylene signaling in regulating carotenoid levels post-RR stage. The retention of carotenoids and high sucrose in *atr-1* highlights that novel alleles in *ETR* genes can be used to improve on-vine-ripened tomato fruit quality. In the future, the targeted mutagenesis of different domains of ETRs may lead to tomato varieties with long shelf life and high nutritional quality.

## Materials and Methods

### Plant Growth Conditions

Tomato (*Solanum lycopersicum*, cv. Ailsa Craig) seeds were surface-sterilized and sown on filter papers. After the emergence of the radicle, seedlings were grown on Soilrite (https://www.keltechenergies.com/soilrite-mix.html) either under continuous white light (100 µmole m^-2^s^-1^) or in darkness at 25°C. Unless otherwise mentioned for all experiments, the plant age was counted from the time point of emergence of the radicle. For all experiments, seeds multiplied from the mutant line *atr-1* that was backcrossed twice were used. Plants were grown in the greenhouse under the natural photoperiod (12-14h day, 28±1°C; 10-12h night, 14-18°C). Fruits were harvested from the first two trusses at different stages of ripening, viz., Mature Green (MG), Breaker (BR), and Red ripe (RR) stages. Fruits were also harvested at RR+7 days (RR+7D) and RR+15 days (RR+15D)

### Mutants Screening and Genetic Analysis

The protocol of EMS mutagenesis and collection of M_2_ seeds was essentially similar to that described in **Al-Hammady et al. (2003).** The M_2_ seeds were germinated on Soilrite in tightly sealed boxes with parafilm, along with calcium carbide (15 gm) encased in a perforated polythene pouch. The calcium carbide, being hygroscopic using humid air inside the sealed box, slowly releases acetylene, leaving residual calcium oxide (lime). The boxes were incubated in darkness (25±2°C) and observed under the green safelight for the seedling triple response. The taller seedlings displaying a lack of triple response were selected as putative mutants and were grown to maturity in the greenhouse. Out of six lines, five perished in the greenhouse, and one that survived was named *a*ce*t*ylene-*r*esistant 1 (*atr-1*) mutant. For the F_2_ segregation analysis, seeds were germinated in the presence of acetylene, and after 12 days, the seedlings were scored for the presence or absence of the triple response.

### Physiological characterization

For growing seedlings in the presence of 1-aminocyclopropane carboxylic acid (1- ACC), ABA, kinetin, glucose, and Norflurazon, seeds were germinated in water-agar (0.8%) supplemented with the respective compound. For ethylene treatment, seedlings were grown in air-tight boxes, and ethylene was injected into the box through a rubber septum (2 mL L^-1^). The 1-methylcyclopropene (1-MCP) treatment and root penetration scoring was carried out in air-tight boxes as described in **Santisree et al. (2011).** For the senescence study, the leaves were harvested from the 8th node of 45-day-old plants and were incubated in air or ethylene (2 mL L^-1^) on moist filter papers in darkness (25±2°C). For monitoring the dark-induced senescence. 15-day-old light-grown seedlings were incubated in total darkness for ten days.

For cotyledon abscission and flower drop assay, 5-day-old light-grown seedlings or detached inflorescence were kept in sealed containers with air or ethylene (2 mL L^-1^). Ethylene was injected into the container through a rubber septum fitted in the lid. For the thermotolerance study, the fruits harvested at RR were kept at 45±2°C for three days. For post-harvest shelf life, RR fruits were kept at room temperature for 30 days, and weight loss was monitored. The internode lengths were measured for at least five to seven successive nodes from three-month-old plants.

### Estimation of ethylene emission

For ethylene emission, seedlings were grown on moist Soilrite in light (100 μmole m^-2^s^-1^). Before ethylene estimation, the boxes were sealed for six hours; after that, ethylene (1 mL) was withdrawn from the headspace with a syringe. Fruits harvested at different stages of ripening were kept in sealed boxes. After six hours, ethylene (1 mL) was withdrawn from the headspace with a syringe. Ethylene was estimated as described by **Kilambi et al. (2013).**

### Biochemical characterizations

The °Brix, fruit diameter, and firmness were determined as described by **Gupta et al. (2014).** The total chlorophyll level in leaves was determined using the method of **Porra et al. (1989)**. The carotenoid profiling was carried out following the procedure of **Gupta et al. (2015).** The phytohormone levels in seedlings and fruits were determined using Orbitrap ExactivePlus LC-MS following a previously described protocol (**Pan et al., 2010; Bodanapu et al., 2016**). The primary metabolite analysis from seedlings and fruits using Leco Pegasus GC-MS was carried out by a protocol modified from **Roessner et al. (2000)** described in **Bodanapu et al. (2016).** Only metabolites and phytohormones with a ≥1.5-fold change (log2 ± 0.584) and p-value ≤0.05 were considered to be significantly different. Principal Component Analysis (PCA) was performed using Metaboanalyst 5.0 (http://www.metaboanalyst.ca/). Heat maps were generated using Morpheus (https://software.broadinstitute.org/morpheus/).

### Gene Expression Analysis

The seedlings or the pericarp from fruit tissue were frozen in liquid N_2_ and homogenized to a fine powder. About 100 mg of homogenized powder was used for total RNA extraction with the RNeasy kit (Qiagen) according to the manufacturer’s instructions. Total RNA was reverse transcribed into cDNA using the first strand cDNA synthesis kit (Invitrogen, India) following the manufacturer’s instructions. PCR on cDNA was performed using the actin gene as control, and the PCR products were analyzed on 1.2 % (w/v) to 2% (w/v) agarose gel based on the product size. The primer sequences are given in **Table S2.**

### Candidate gene analysis

The cDNA was prepared from mRNA isolated from leaves of 15-day-old plants using an Invitrogen cDNA synthesis kit. As candidate genes, *LeEIN2*, *LeEIL1*, *LeEIL2*, and *LeEIL3* were examined for any mutation. The primer sequences are given in **Table S3.** The overlapping sets of primers were designed to cover the full gene length. The presence of the mutation in the genes was examined by the CEL-I endonuclease assay (**Mohan et al., 2016**).

### Genome Sequencing

Whole Genome Sequencing and data analysis of Ailsa Craig and *atr-1* were done as described in **Gupta et al. (2022).** Briefly, sequencing was performed on the HiSeqX sequencing system (Illumina) (GeneWiz Inc. NJ, USA) according to the manufacturer’s protocol. A total of ∼200 million reads (31.5 Gb) with Q30 > 29 Gb data was generated. The raw reads were filtered using fastp software (v0.19.5) using parameters -M 30 -3 -5. The 2X 150 bp reads were mapped on *S. lycopersicum* cv. Heinz version SL3.0 using BWA-MEM (0.7.17) (**Li and Durbin, 2009**). The data analysis and variant calling was performed as described by **Gupta et al. (2020).** The resulting vcf (variant calling format) files were annotated using the SIFT4G algorithm. The effect of base substitutions on protein function was determined by SIFT4G (SIFT score ≤0.05 is considered deleterious) using the SIFT4G- ITAG3.2 genome reference database generated by **Gupta et al. (2020).** The modeling of WT and ETR4 protein was carried out using DDmut software (https://biosig.lab.uq.edu.au/ddmut).

### Statistical analysis

A minimum of three independent biological replicates was used for all experiments. Statistical Analysis On Microsoft Excel (https://prime.psc.riken.jp/MetabolomicsSoftware/StatisticalAnalysisOn-MicrosoftExcel) was used to obtain significant differences between data points. A Student’s *t*-test was also performed to determine significant differences (* for P ≤0.05).

## Data Availability Statement

All data pertaining to this study are included within the article and its supplementary files.

## Author Contributions

YS and RS designed this project and wrote the manuscript. SKG and PS performed most of the experiments. PG analyzed the AC and *atr-1* genome sequences. HVK did fruit RT-PCR analysis. All authors read and approved the manuscript.

