## Supplementary material for "A tomato ethylene-insensitive mutant displays altered growth and higher β-carotene levels in fruit": Figure S

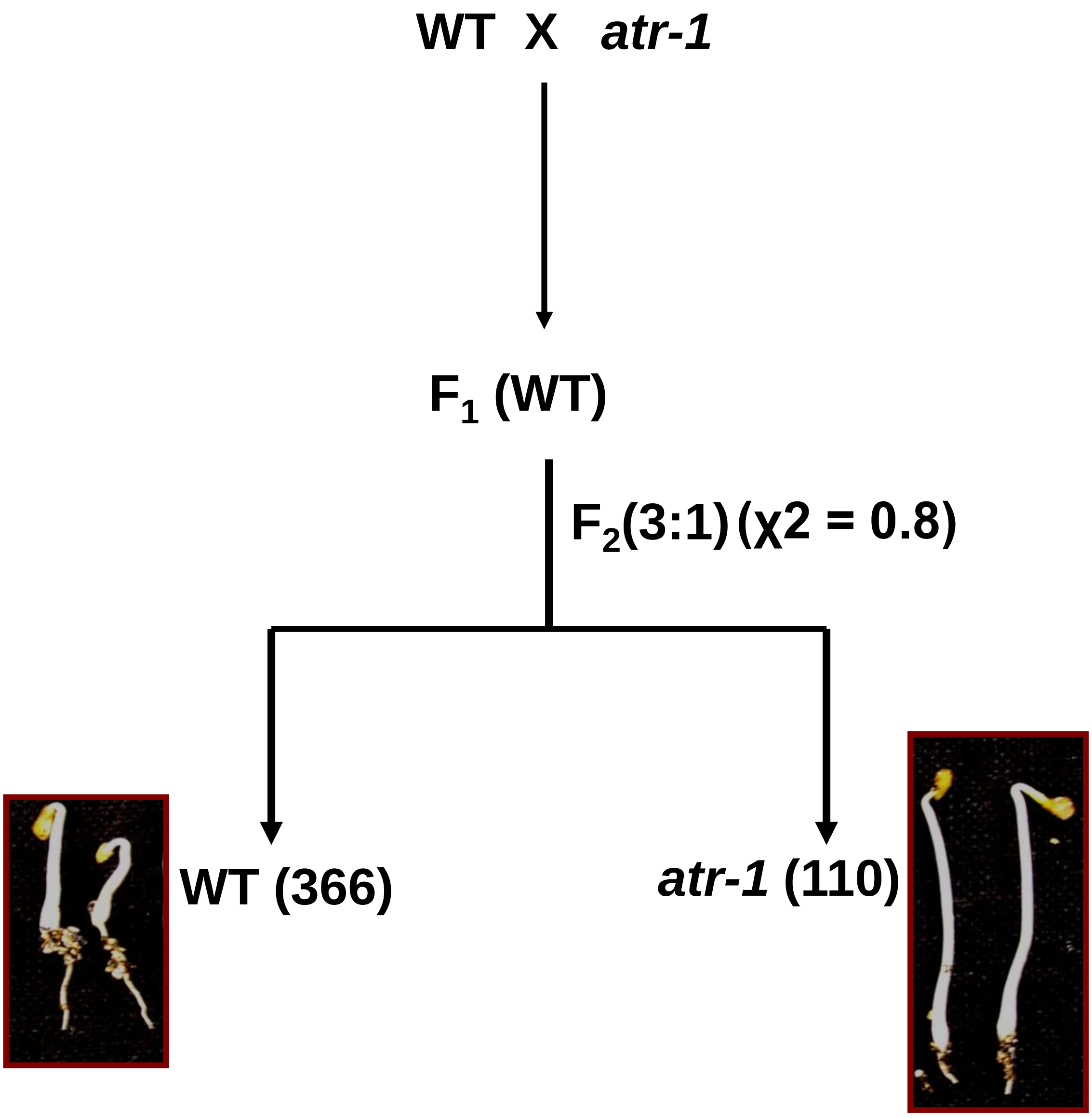


**Figure S1**. Genetic Segregation of *atr-1* mutant after crossing with WT. The $F_2$ seeds were germinated in darkness in the presence of acetylene, and seedlings were scored based on the presence or absence of triple response phenotype. The numbers in parentheses indicate the number of seedlings showing triple response or insensitive phenotypes.

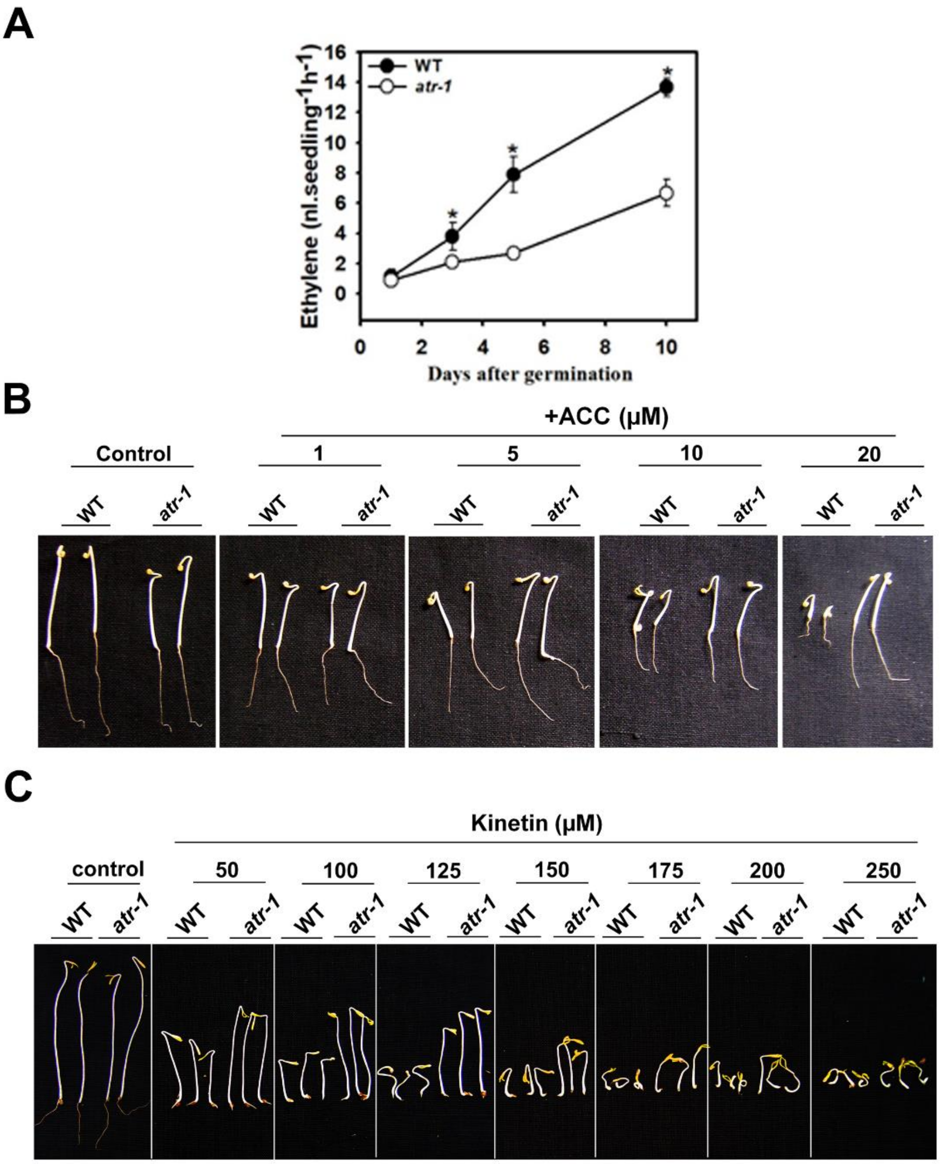


**Figure S2**. Characterization of dark-grown seedlings of *atr-1*. (**A**) Ethylene emission from dark-grown *atr-1* and WT seedlings. Asterisks indicate statistically significant differences between WT and *atr-1* seedlings. (Student's t-test, * < 0.05). (**B**) WT and *atr-1* seedlings were grown in darkness with increasing concentrations of 1-aminocyclopropane carboxylic acid (ACC). Seedlings were photographed after five days of ACC treatment. Each experiment contained 7 to 8 seedlings and was repeated more than three times. (**C**) Effect of varying concentrations of kinetin on the growth of etiolated seedlings of WT and *atr-1*. The photographs show 7-day-old seedlings. Each experiment contained 40-45 seedlings and was repeated more than three times.

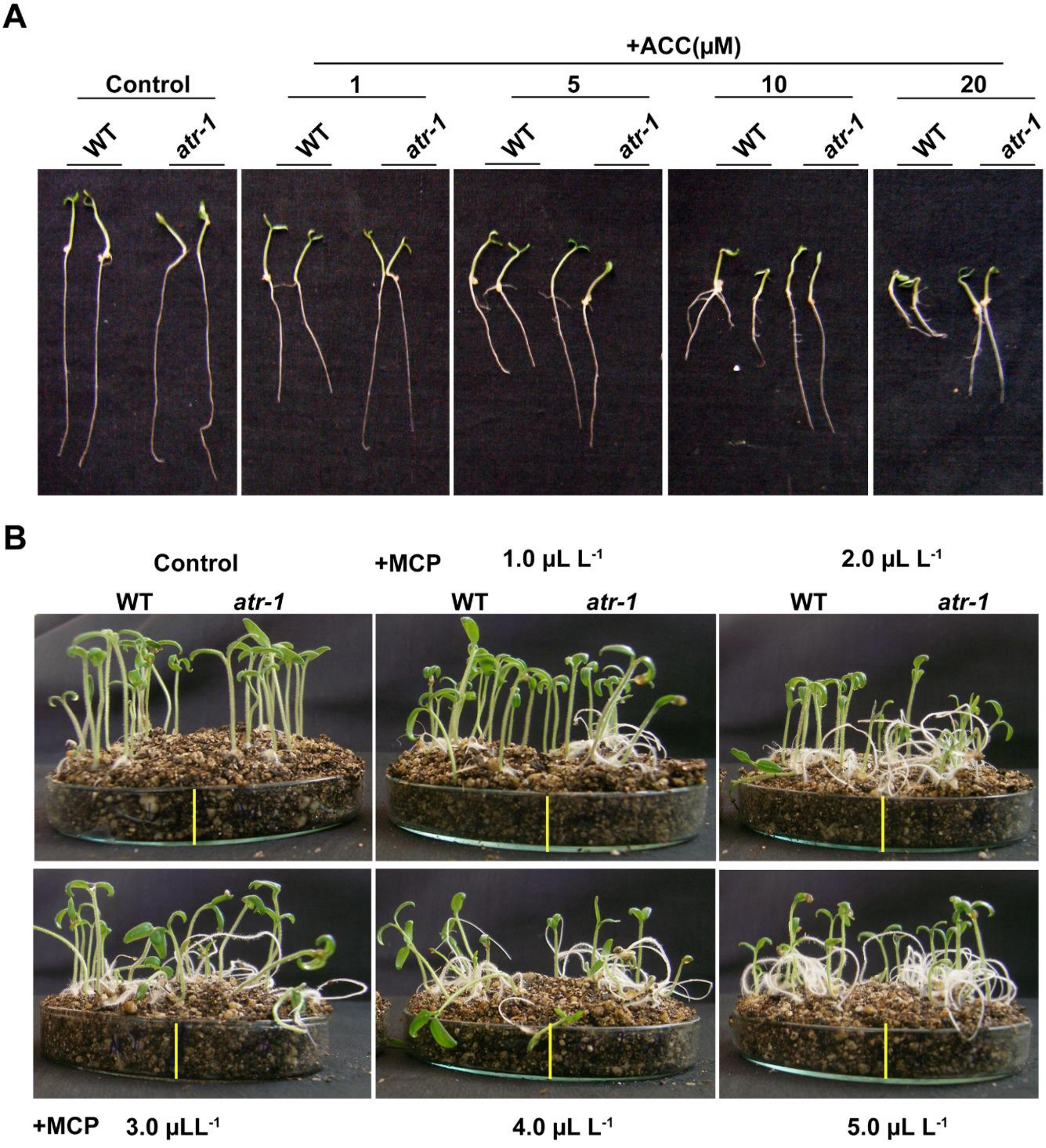


**Figure S3.** Effect of ACC and 1-MCP on the growth of light-grown *atr-1* seedlings. (**A**) 7-day-old light-grown WT and *atr-1* seedlings grown in different concentrations of ACC. The experiment was repeated three times, and each replicate consisted of 10-12 seedlings. (**B**) Root penetration in *atr-1* mutant is more sensitive to 1-MCP. Photographs show the effect of increasing 1-methylcyclopropane (1-MCP) concentration on root penetration of 7-day-old light-grown WT and *atr-1* seedlings. The vertical yellow line on the photograph marks the separation of WT (left) and *atr-1* (right) seedlings. The experiment was repeated two times, and each replicate consisted of 10-12 seedlings.

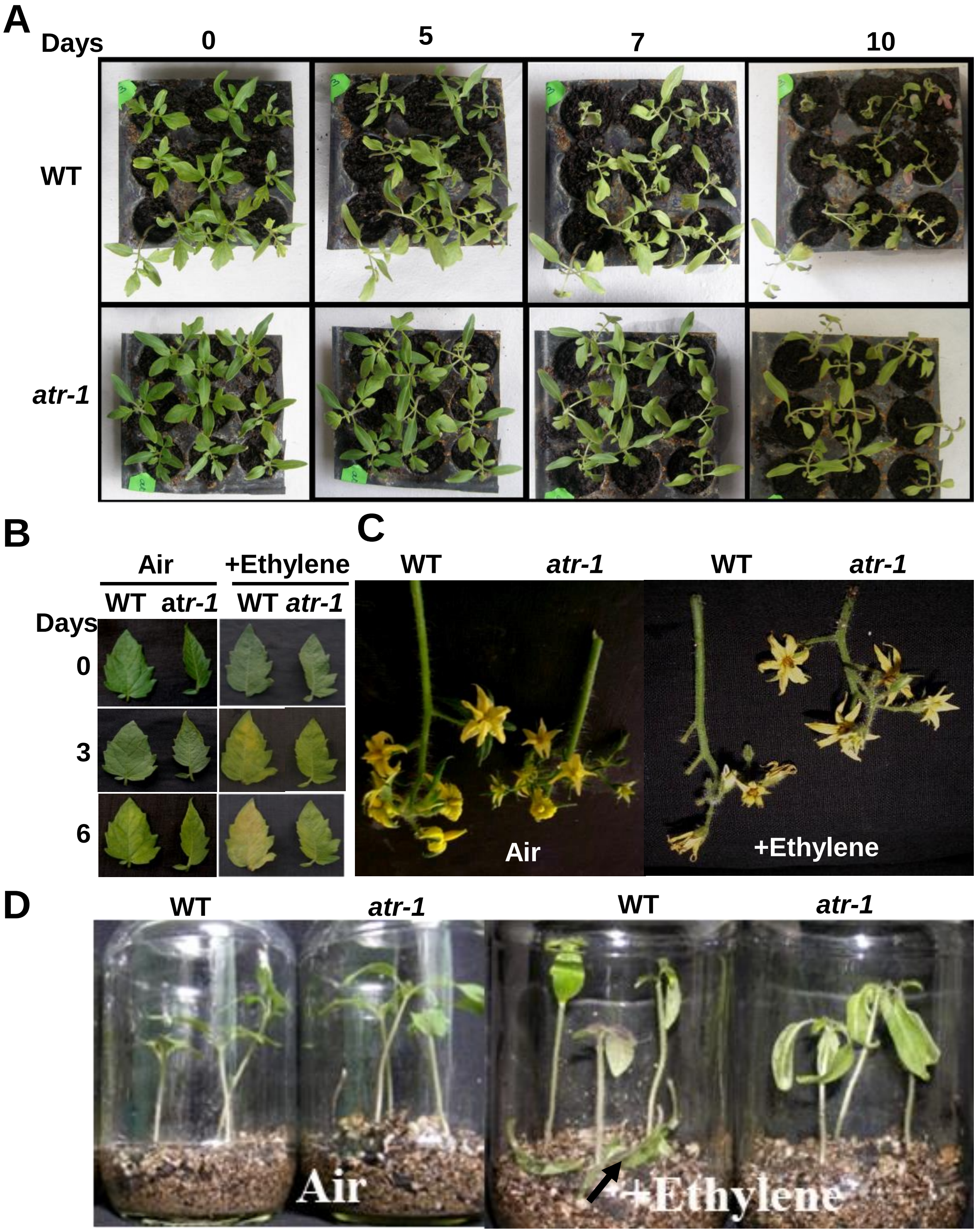


**Figure S4.** Effect of ethylene on senescence and abscission in *atr-1*. (**A**) The *atr-1* mutant shows sluggish dark-induced senescence. The 15-day-old light-grown *atr-1* and wild-type (WT) seedlings were incubated in total darkness for 10 days. Note: For taking photographs, the seedlings were briefly taken out to natural light and were transferred back to darkness. The experiments were repeated more than three times, and in each replicate consisted of 8-10 seedlings. (**B**) Effect of *atr-1* mutation on leaf senescence. The detached leaflets from the 8$^{th}$ node of WT (left) and mutant (right) were treated with air or 2 mL $L^{-1}$ ethylene for 6 days in darkness. The leaflets were photographed at different time points, indicated on the left of each panel. (**C**) The picture shows the inflorescence of WT and *atr-1* after 24 h of exposure to 2 mL $L^{-1}$ ethylene (right) and without ethylene (left). **(D)** Cotyledon abscission in WT and *atr1* plants. Two-week-old plants were sealed in airtight chambers for 24 h in the absence (air; left) and presence of ethylene (right; 2 mL $L^{-1}$). The black arrow points to abscised cotyledons lying on the Soilrite.

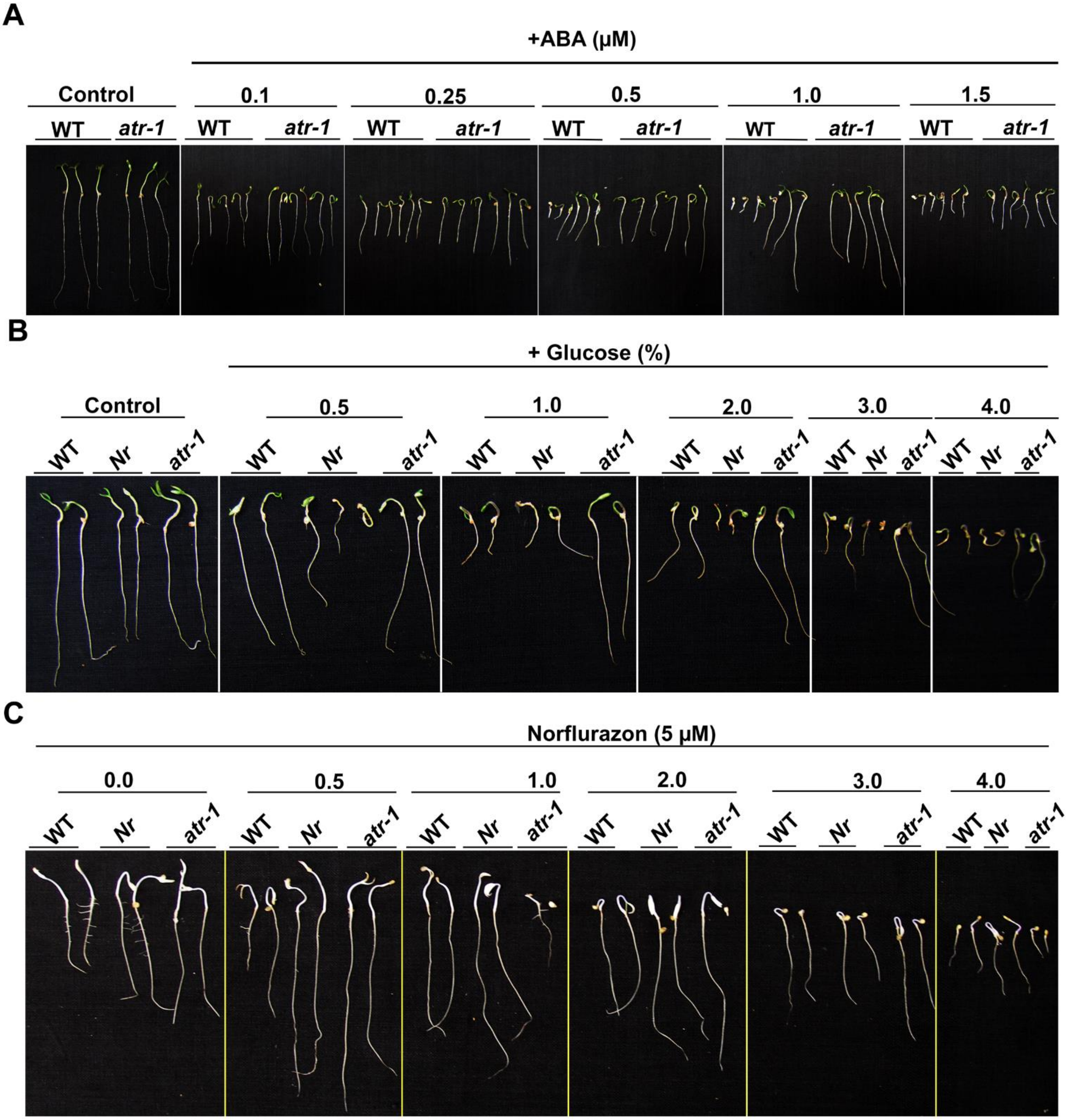


**Figure S5.** The effect of ABA, glucose, and Norflurazon on the growth of *atr-1* and WT seedlings. (**A**) The effect of ABA on hypocotyl and root growth of light-grown *atr-1* and WT seedlings. The seedlings were photographed after eight days from germination. (**B**) Effect of glucose on growth of 8-day-old WT, *Nr,* and *atr-1* seedlings grown in different glucose concentrations. (**C**) Effect of Norflurazon (an ABA inhibitor) on growth on 8-day-old WT, *Nr,* and *atr-1* seedlings. Note that Norflurazon relieves the inhibitory effect of glucose on WT, *Nr,* and *atr-1* seedling development. Photographs show 8-day-old WT, *Nr,* and *atr-1* seedlings grown in different concentrations of glucose (%)) supplemented with five µM Norflurazon. The experiment was repeated more than three times for each treatment, and each replicate consisted of 10-12 seedlings.

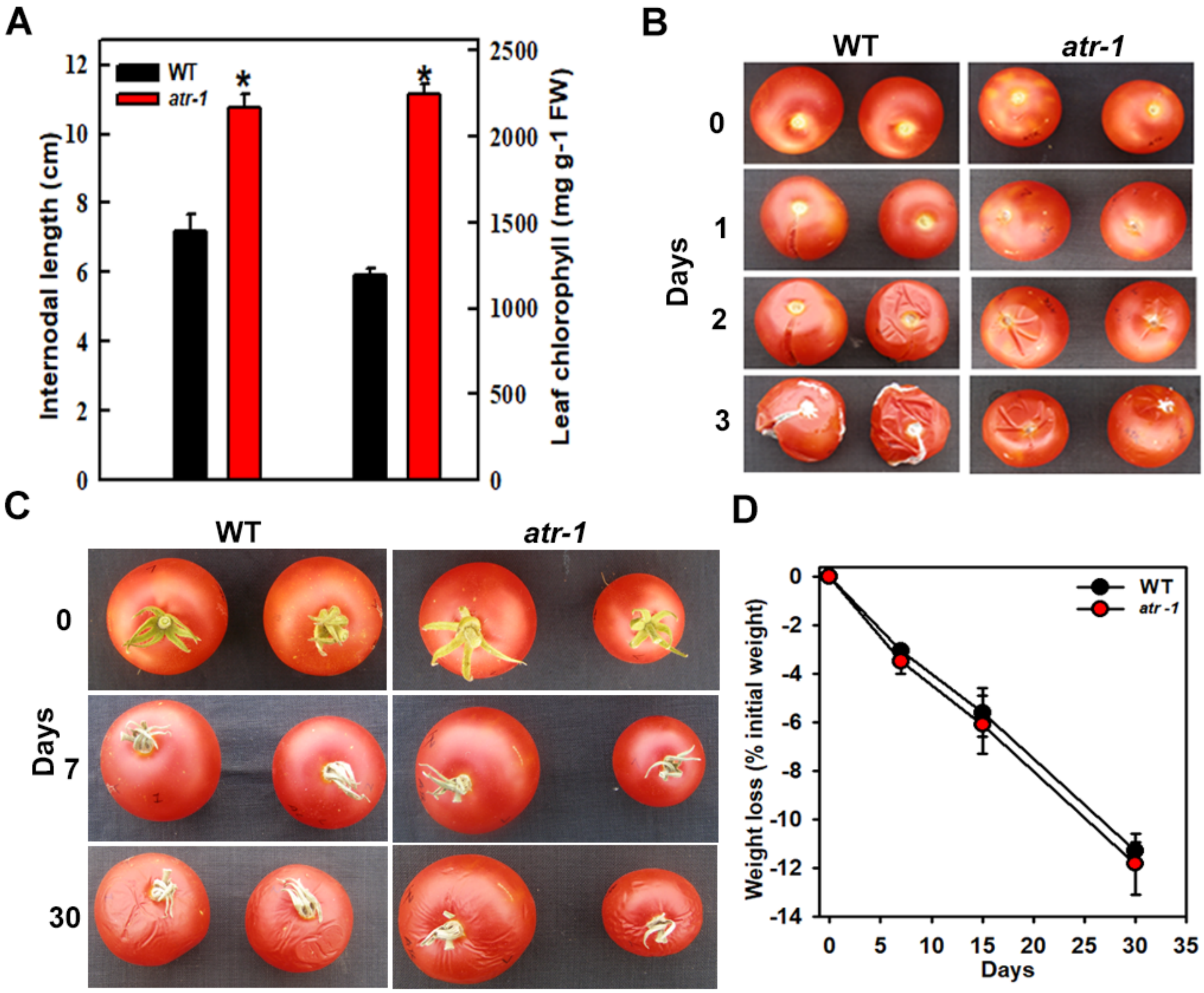


**Figure S6:** Characterization of vegetative growth, thermotolerance, and post-harvest shelf life of *atr-1*.(**A**) Internode length of WT and *atr-1* mutant (Left). Five measurements were taken from the sixth node of five 3-month-old WT and *atr-1* mutant plants. Quantification of total leaf chlorophyll content in leaflets from the eighth node of WT and *atr-1* mutant plants (Right). Asterisks indicate statistically significant differences between WT and *atr-1*. (Student's t-test, * < 0.05). (**B**) The WT and *atr-1* fruits after harvest at the red ripe stage were incubated at 45°C. Fruits were photographed on different days indicated on the left of respective photo panels. *atr-1* fruits were more resistant to heat, as visualized by delayed fruit skin cracking (**C**). Red ripe fruits of *atr-1* and WT incubated under continuous white light at room temperature. Fruits were photographed on different days indicated on the left of respective photo panels. The post-harvest shelf life of fruits was nearly the same, as monitored by the appearance of wrinkles on fruits. (**D**) Loss of water from *atr-1* and WT fruit harvested at RR stage and shown in panel (**C**). The experiment was repeated five times, and each replicate consisted of 5 fruits.

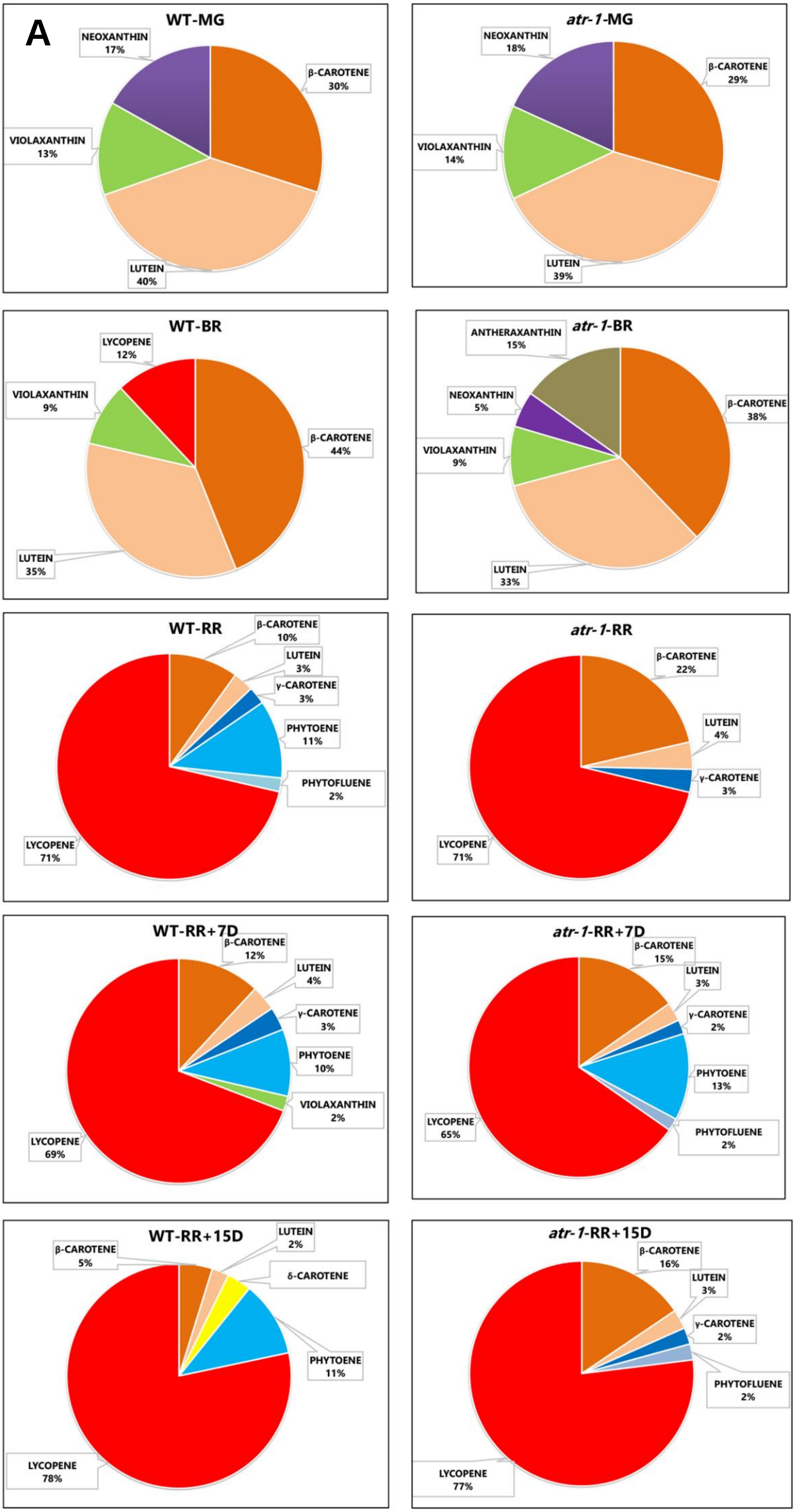
A
WT-MG
NEOXANTHIN 17%
β-CAROTENE 30%
VIOLAXANTHIN 13%
LUTEIN 40%
atr-1-MG
NEOXANTHIN 18%
β-CAROTENE 29%
VIOLAXANTHIN 14%
LUTEIN 39%
WT-BR
LYCOPENE 12%
VIOLAXANTHIN 9%
β-CAROTENE 44%
LUTEIN 35%
atr-1-BR
ANTHERAXANTHIN 15%
NEOXANTHIN 5%
β-CAROTENE 38%
VIOLAXANTHIN 9%
LUTEIN 33%
WT-RR
β-CAROTENE 10%
LUTEIN 3%
γ-CAROTENE 3%
PHYTOENE 11%
PHYTOFLUENE 2%
LYCOPENE 71%
atr-1-RR
β-CAROTENE 22%
LUTEIN 4%
γ-CAROTENE 3%
LYCOPENE 71%
WT-RR+7D
β-CAROTENE 12%
LUTEIN 4%
γ-CAROTENE 3%
PHYTOENE 10%
VIOLAXANTHIN 2%
LYCOPENE 69%
atr-1-RR+7D
β-CAROTENE 15%
LUTEIN 3%
γ-CAROTENE 2%
PHYTOENE 13%
LYCOPENE 65%
PHYTOFLUENE 2%
WT-RR+15D
LUTEIN 2%
β-CAROTENE 5%
δ-CAROTENE
PHYTOENE 11%
LYCOPENE 78%
atr-1-RR+15D
β-CAROTENE 16%
LUTEIN 3%
γ-CAROTENE 2%
PHYTOFLUENE 2%
LYCOPENE 77%

Figure S7A

**B**

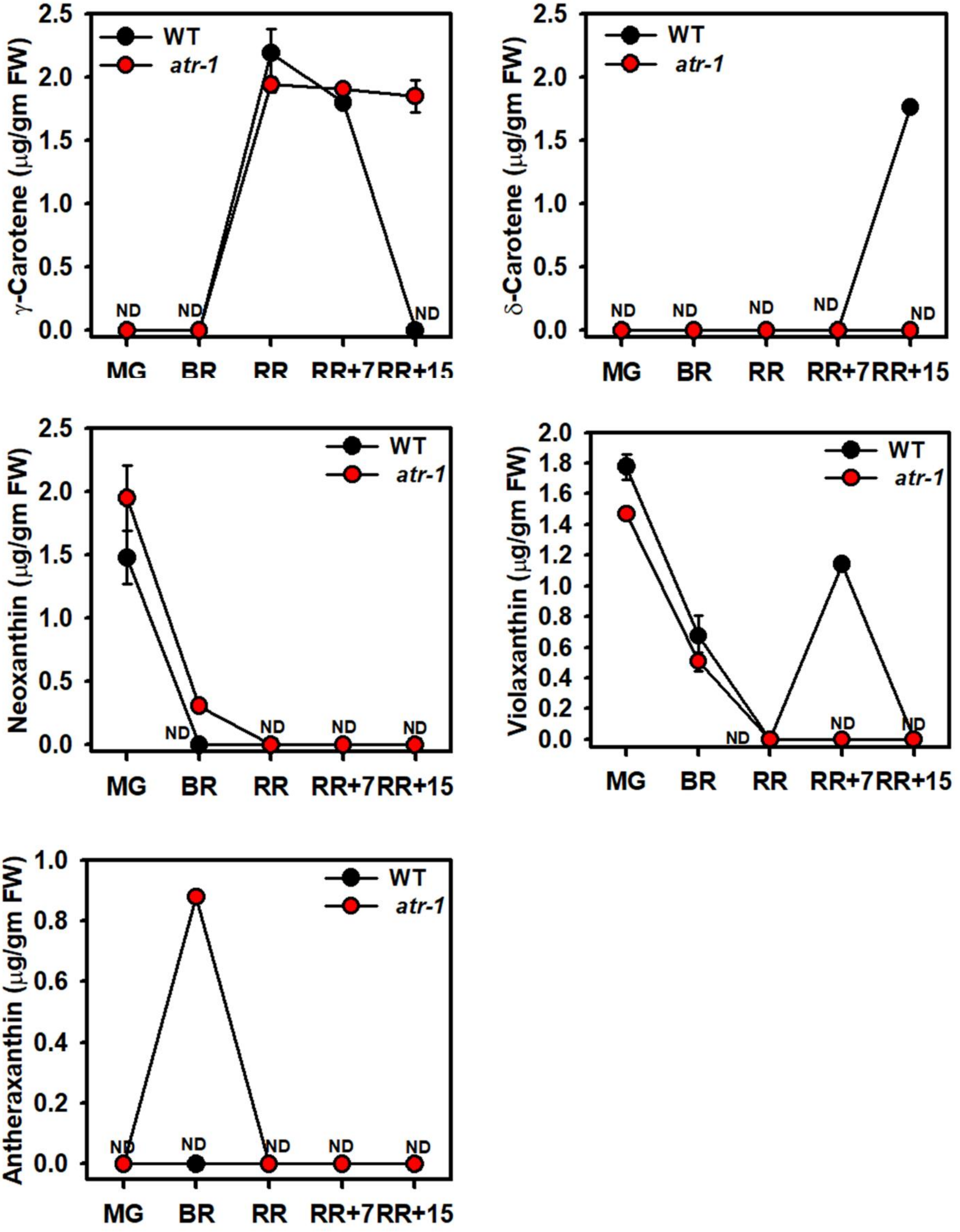


**Figure S7.** The distribution of carotenoids in *atr-1* and WT fruits. (**A**) Pie diagrams show relative levels of carotenoids at different ripening stages. Note high β-carotene levels in *atr-1* at RR and later stages. (**B**) Change in different carotenoid levels during the ripening in *atr-1* and WT (AC) fruits. The carotenoid data are expressed as mean SE ($n \geq 3$). *$P \leq 0.05$. **See Dataset S3** for individual carotenoid levels and significance. ND-not detected

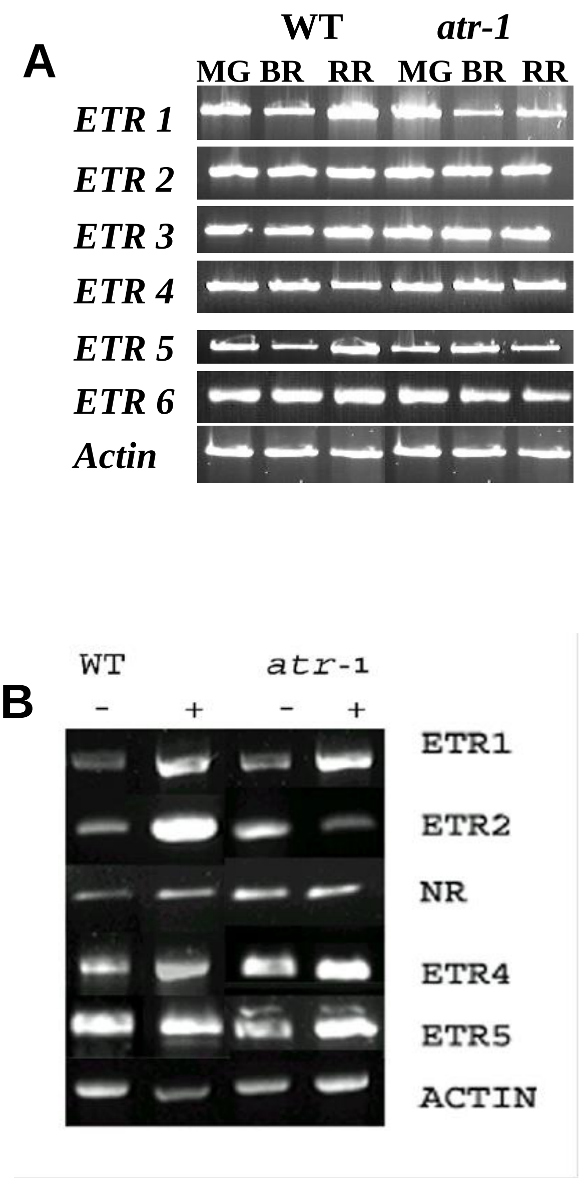


**Figure S8.** Expression of ETR genes in WT and *atr-1* mutant. **A.** Expression of *ETR* genes in WT and *atr-1* mutant at different stages of fruit ripening. Expression of the actin gene was used as an internal control to ensure equal loading of the samples. **B.** Expression of *ETR* genes in WT and *atr-1* mutant 12-day-old etiolated seedlings grown with (+) or without acetylene (-). Expression of the actin gene was used as an internal control to ensure equal loading of the samples. For primer sequences, see **Table S2.**

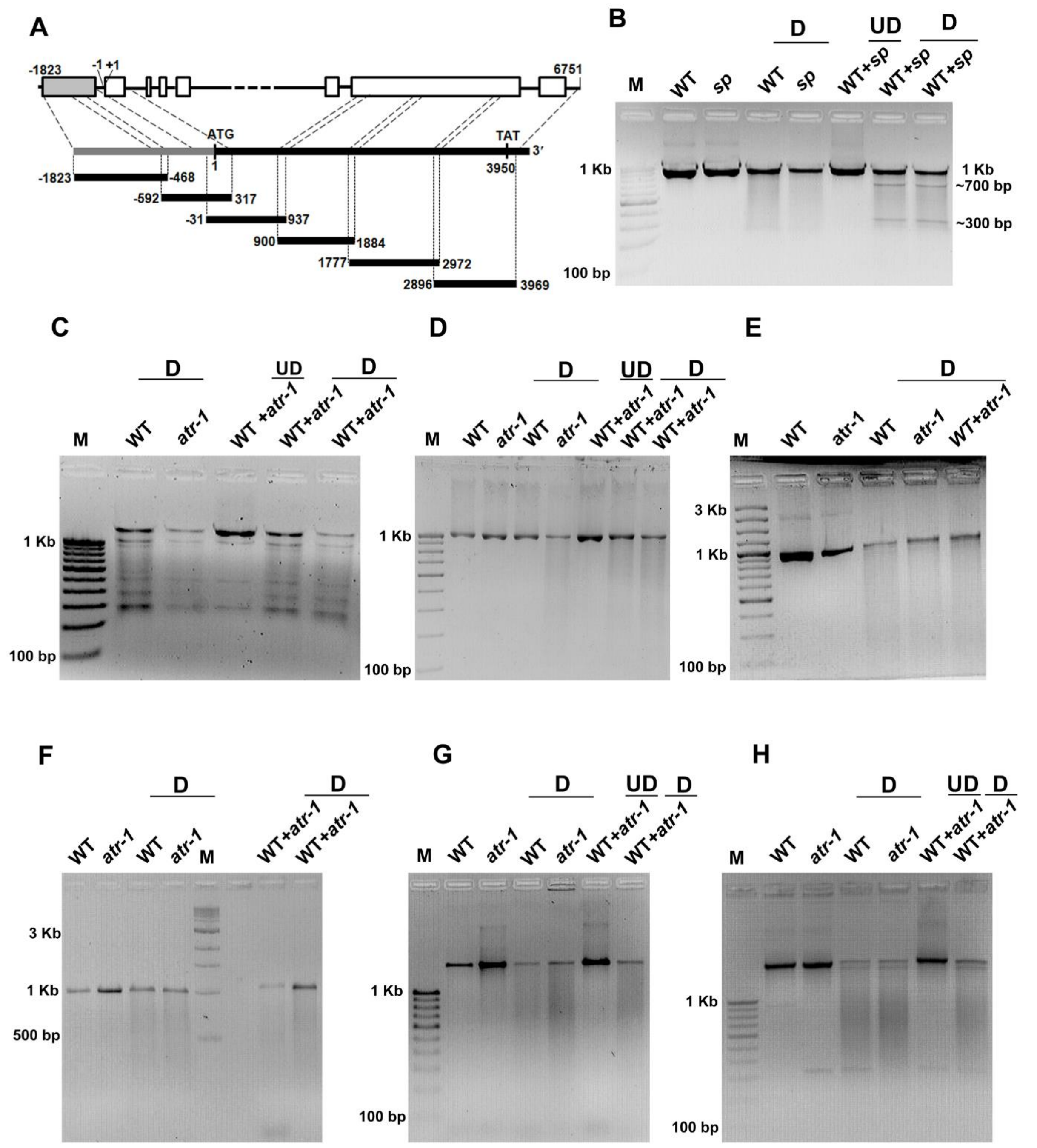


**Figure S9.** Analysis of mutation in tomato *EIN2* gene. (**A**) The gene structure of the *EIN2* gene. A thick line indicates the six introns, boxes represent seven exons, and the gray box represents the promoter region.  The six sets of overlapping primers were designed from the exons and promoter region of the gene to cover the promoter and entire CDS sequences. The dotted lines represent the primer position in CDS sequences, and the dashed line represents its position on genomic coordinates. (**B**) CEL I digestion profile of *sp* mutants, which was used as a positive control. The heteroduplex lane WT+*sp* show the presence of two digested band, whereas it is not present in the homoduplex lane of either WT or *sp* mutant. (**C-H**) CEL I digestion of *SlEIN2* amplified product from the promoter region and cDNA of WT and *atr-1* mutant. No digested fragments were observed in the heteroduplex or homoduplex lane of any sets of CEL I digestion, indicating that the *atr-1* mutation is not located in the *EIN2* gene. **D**= digested, **UD**= undigested. **M**= marker (size in bp). For primer sequences, see **Table S3.**

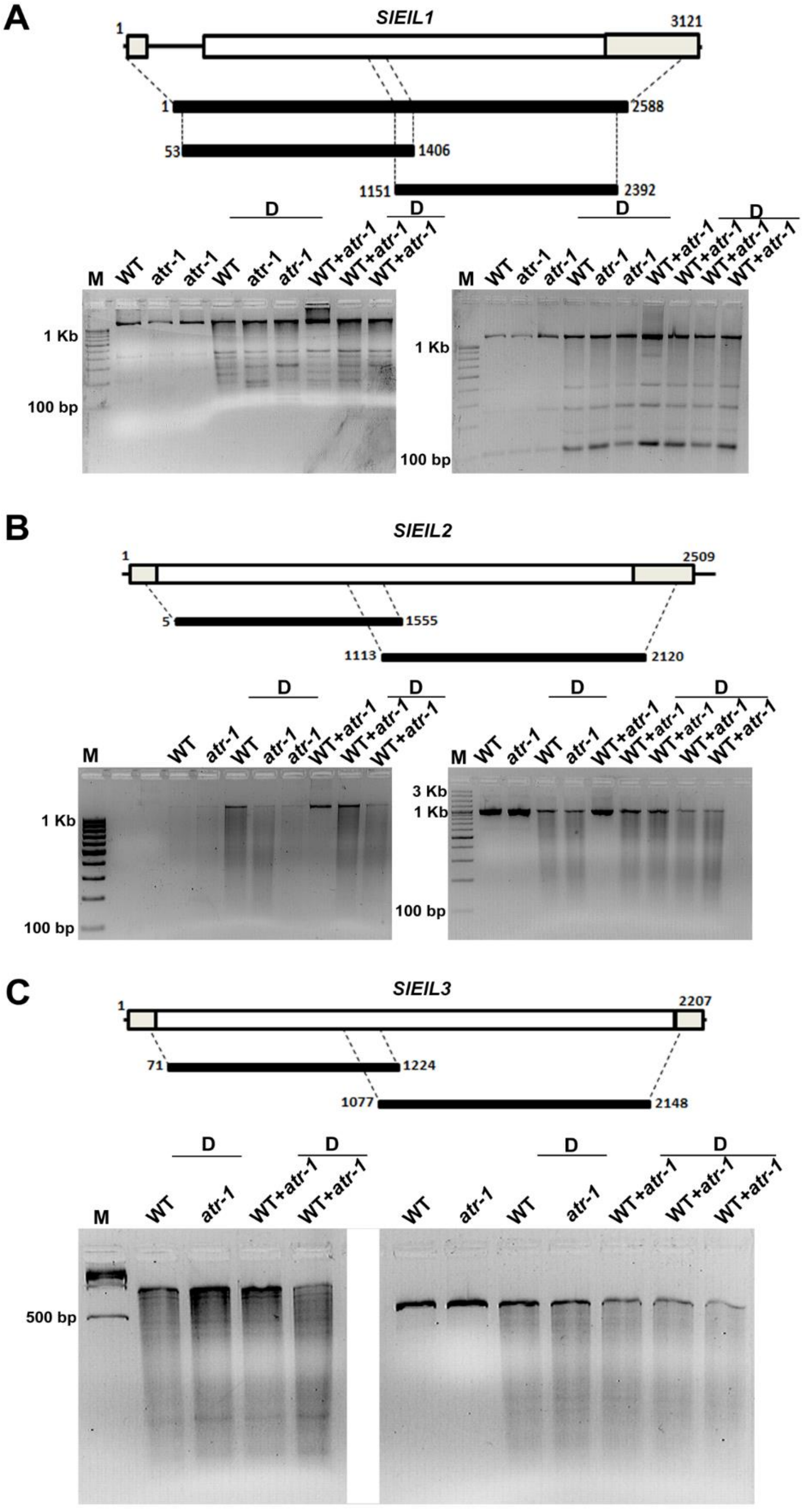


**Figure S10.** Mismatch cleavage analysis of tomato *EIL* genes in *atr-1* mutant. (**A**) The gene structure of the *EIL1* gene. Lines indicate the introns, the CDS region is represented by boxes, and the gray color box represents the UTRs region. The primers were designed from mRNA sequences. The dotted lines represent the primer position on cDNA, and the dashed lines represent its position on genomic coordinates. (**B, C**) *EIL2* and *EIL3* gene structure and positions of primers. Both genes consist of a single exon and thus have no intron. Two primer sets were designed to cover the complete sequences of CDS at the positions marked by dotted lines. The *EIL1*, *EIL2*, and *EIL3* genes from both WT and *atr-1* amplified with both sets of primers were digested by CEL-I. The gel images are shown below the respective genes. The left gel image is from primer set 1, and the right is from primer set 2. No mutation was observed in the *EIL1*, *EIL2*, and *EIL3* genes of the *atr-1* mutant. For primer sequences, see **Table S3.**

**A**

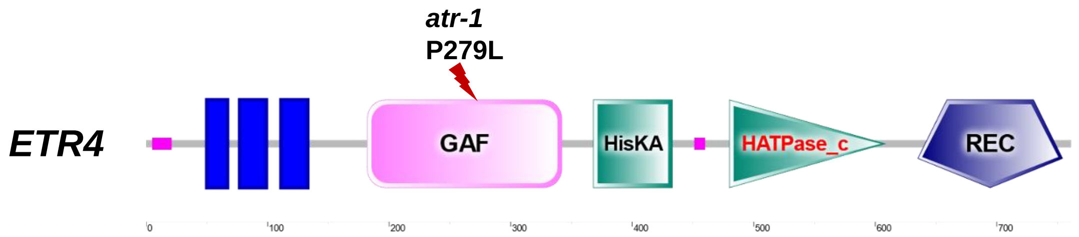


**B**

MLRTLASALLVLSFFVSLSAADNGFPRCNCDDEGFWSIESILECQKISDLFIAIAYFSIP

IELLYFVSCSNFPFKWVLFQFIAFIVLCGMTHLLNFWTYYGQHPFQLMLALTIFKVLTAL

VSFATAITLITLFPMLLKVKVREFMLKKKTWDLGREVGLIKMQKEAGWHVRMLTQEIRKS

LDRHTILYTTLVELSKTLDLHNCAVWKPNENKTEMNLIHELRDSSFNSAYNLPIPRSDPD

L

VIQVKESDGVKILDADSPLAVASSGGSREPGAVAAIRMPMLKVSNFKGGTPELVPECYAI

LVLVLPSEQGRSWCSQEIEIVRVVADQVAVALSHAAILEESQHMRETLEEQNRALEQAKQ

DALRASQARNAFQMVMSHGLRRPMHSILGLLSLLQDEKLGNEQRLLVDSMVKTSNVVSTL

IDDVMDTSTKDNGRFPLEMRYFQLHSMIKEAACLAKCLCAYRGYNISIEVDKSLPNHVLG

DERRVFQVILHMVGNLLKDPNGGLLTFRVLPESVSREGIGGAWRTRRSNSSRDNAYIRFE

VGTSNNHSQPEGTMLPHYRPKRCSKEMDEGLSFTVCRKLVQLMQGDIWVIPNPEGFDQSM

AVVLGLQLRPSIAIGIPEYGESSDHSHPHSLLQGVKVLLADYDDVNRAVTSKLLEKLGCS

VSAVSSGRDCIGVLSPAVSSFQIVLLDLHLPDLDGFEVTMRIRKFGSHNWPLIVGLTATA

DENVTGRCLQIGMNGLIRKPVLLPGIADELQRVLLRGSRMM

**Figure S11.** The protein sequence and different domains of tomato ETR4. **(A)** The SMART (http://smart.embl-heidelberg.de/) images ETR4 protein showing the location of the mutation in the GAF domain in *atr-1*. The transmembrane domain, GAF domain, histidine kinase A domain, histidine kinase-like ATPase, and receiver domains are shown in the cartoon. (**B**) The amino acid sequence of the tomato ethylene receptor ETR4. The upward arrow points to the location of the mutation P279L.

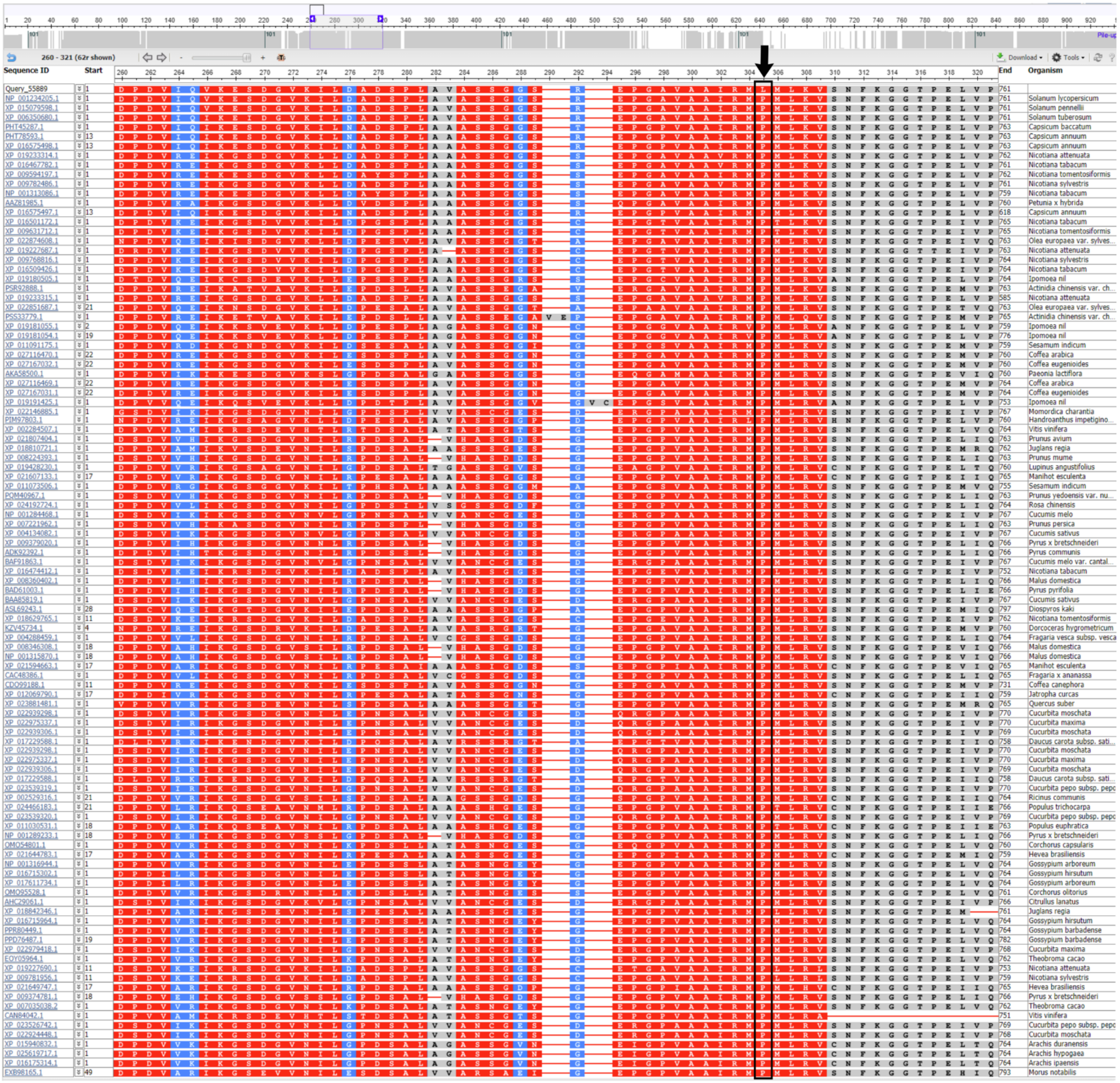

**Figure S12.** Alignment of the GAF domain of different species. Substitution of amino acid P at 279 by amino acid L in the GAF domain of ETR4 protein of *atr-1* mutant, used as a query sequence. The amino acid P (enclosed in a square box and indicated by an arrow) is highly conserved in closely related 100 plant species.

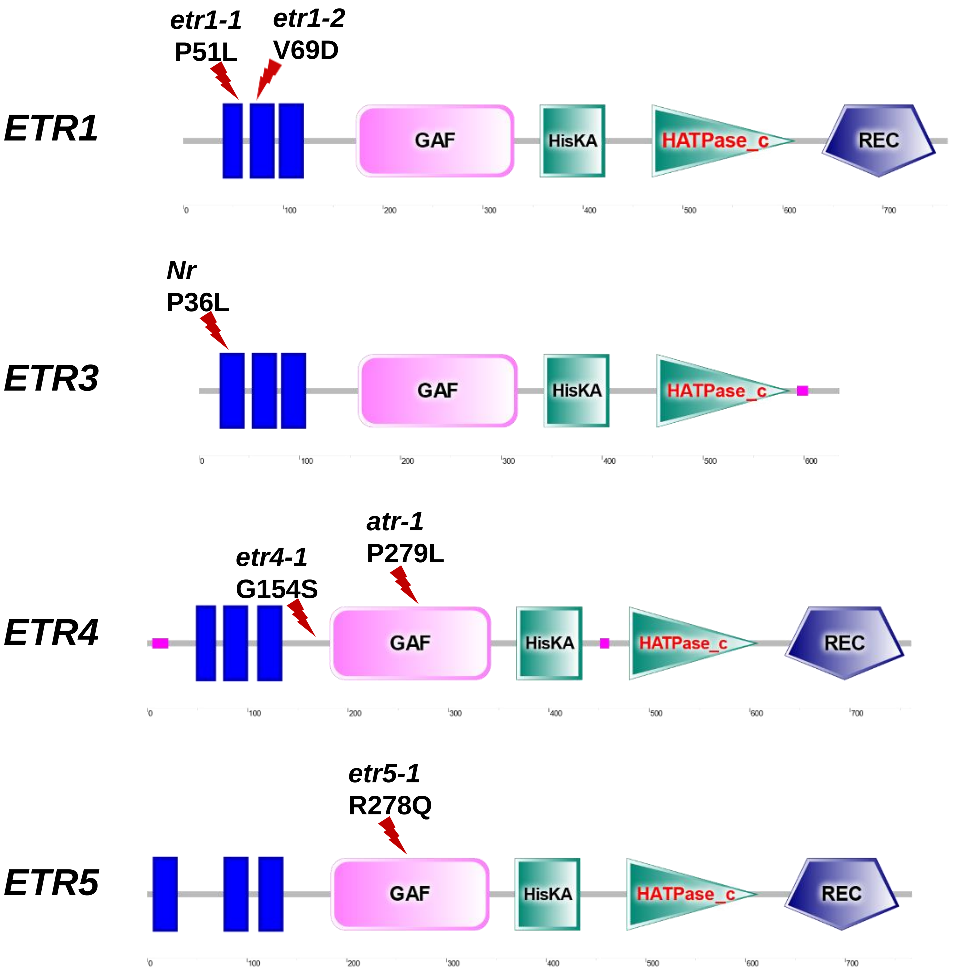


**Figure S13**. Location of mutant site in tomato ETR mutants. The SMART (http://smart.embl-heidelberg.de/) images of the different tomato ETR mutants, including *atr-1*. The location of the mutation, associated amino acid change, and the mutant IDs are marked on respective ETR. The transmembrane domain, GAF domain, histidine kinase A domain, histidine kinase-like ATPase, and receiver domains are shown in the cartoons. For phenotypes affected and references, see **Table S1.**

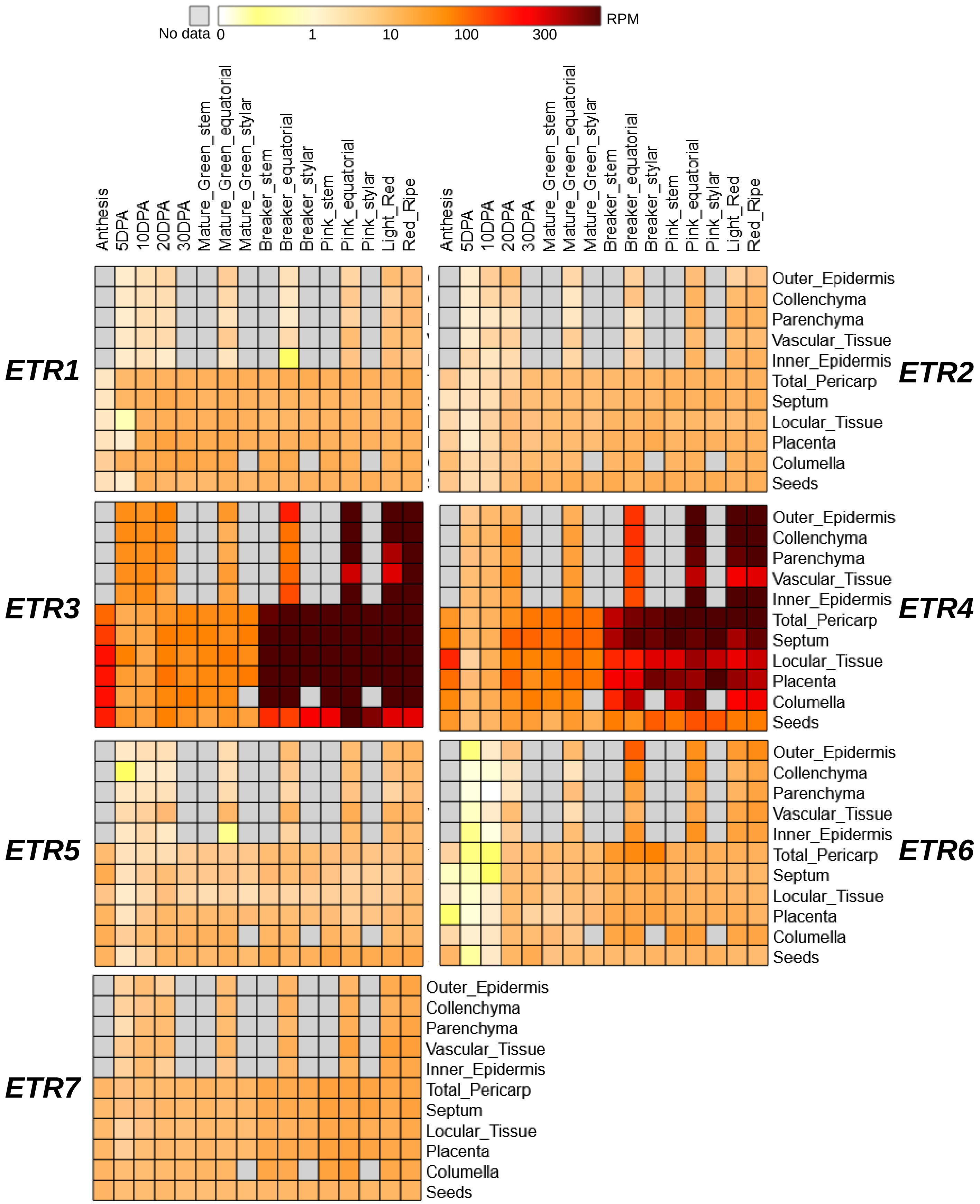


**Figure S14**: The expression of *ETR1-7* genes in tomato fruits from anthesis to full ripening (Data source https://tea.solgenomics.net/expression_viewer/output). The top panel shows the colors used in the heat map for different rpm values. Note the highly increased expression of *ETR3* and *ETR4* genes after the onset of the fruit ripening.
