## Supplementary material for "A tomato ethylene-insensitive mutant displays altered growth and higher β-carotene levels in fruit": Table S1

**Table S1.** List of EMS-induced or spontaneous tomato *ETR* mutants. The table shows the site of mutation and their influence on plant phenotype and fruit ripening.

| **Mutant** | **Gene and SOL id** | **Amino acid substitution** | **Domain** | **Dominant/ Recessive** | **Ethylene sensitivity** | **Plant appearance** | **Leaf shape** | **Fruit color** | **Post-harvest shelf life** | **Reference** |
| --- | --- | --- | --- | --- | --- | --- | --- | --- | --- | --- |
| *etr1-1* | *ETR1*  *Solyc12g011330* | P51L | First transmembrane domain | Dominant | Completely ethylene insensitive | Like WT | Like WT | Yellow to Orange | Longer | Okabe et al., 2011;  Mubarok et al., 2015 |
| *etr1-2* | *ETR1 Solyc12g011330* | V69D | Second transmembrane domain | Dominant | Moderate ethylene insensitivity | Like WT | Like WT | Red Light | Longer | Okabe et al., 2011;  Mubarok et al., 2015 Lanahan et al 1994 |
| *Nr* | *ETR3 Solyc09g075440* | P36L | First transmembrane domain | Dominant | Ethylene insensitive | Tall | Like WT | Yellow | Longer | Wilkinson et al., 1995; Nascimento et al 2021 |
| *atr-1* | *ETR4 Solyc06g053710* | P279L | GAF domain | Recessive | Reduced ethylene sensitivity | Tall | Bigger leaf than WT | Red | No change | This study |
| *etr4-1* | *ETR4* *Solyc06g053710* | G154S | Between the transmembrane and  GAF domain | Recessive | Low ethylene insensitive | Like WT | Like WT | Red | Slightly longer | Mubarok  et al., 2019 |
| *etr5-1* | *ETR5* *Solyc11g00618* | R278Q | GAF  domain | Recessive | Increased ethylene  sensitivity | Like WT | Like WT | Red | Shorter | Mubarok et al.,  2019 |
