## Supplementary material for "A tomato ethylene-insensitive mutant displays altered growth and higher β-carotene levels in fruit": Table S2

**Table S2:** List of ethylene receptors genes and the primers used for RT-PCR analysis.

| **Organ** | **Gene** | **Primer** | **Sequence (5′🡪3′)** |
| --- | --- | --- | --- |
| Seedlings | *SlETR1* Solyc12g011330 | F | GTGGATTATGGATGCCAACAG |
|  |  | R | TCCAAGACATCGTTGATGAGC |
|  | *SlETR2* Solyc07g056580 | F | AAGGCAGTGTGTCAGTTTCTGC |
|  |  | R | ACATCGCACCCTAGATGCAC |
|  | *SlETR3* Solyc09g075440 | F | TGAGGCTTCAGTTGCCAAAC |
|  |  | R | CATCCCACCATCATCTCCAC |
|  | *SlETR4* Solyc06g053710 | F | ACCCCAATGGAGGTCTTCTC |
|  |  | R | CCTTGGAGGAGTGAGTGTGG |
|  | *SlETR5* Solyc11g006180 | F | TAATCAGGTGATGGGCGATG |
|  |  | R | GAAATCGGTTGCTCCAAAGG |
| Fruits | *SlETR1* Solyc12g011330 | F | TGACCCACAGTTGCCTGCTG |
|  |  | R | TGAGCGTTGTAAAAGGTTGC |
|  | *SlETR2* Solyc07g056580 | F | ACAAGGACTGTGGCAATGGTG |
|  |  | R | GCATGAGCCGTTTTTCATCTCC |
|  | *SlETR3* Solyc09g075440 | F | TGGATGTAGCTCGACAAGAAGC |
|  |  | R | TTTGATAGCGTTGAGCATTCAC |
|  | *SlETR4* Solyc06g053710 | F | ACCCCAATGGAGGTCTTCTC |
|  |  | R | CCTTGGAGGAGTGAGTGTGG- |
|  | *SlETR5* Solyc11g006180 | F | TAATCAGGTGATGGGCGATG |
|  |  | R | GAAATCGGTTGCTCCAAAGG |
|  | *SlETR6* Solyc09g089610 | F | TGCTGTTTGCAGGAAGTTGG |
|  |  | R | CTGAAGCTCATCGGCGAGTC |
| Seedlings/fruits | *Actin* | F | CCAAAAGCCAATCGAGAGAA |
|  |  | R | GGTACCACCACTGAGGACGA |
