## Supplementary material for "A tomato ethylene-insensitive mutant displays altered growth and higher β-carotene levels in fruit": Table S3

**Table S3**. Genes and primers used in candidate gene analysis of *atr-1* mutant.

| **Gene** | **Primer** | | **Start-End position** | **Length** | **Sequence (5′🡪3′)** |
| --- | --- | --- | --- | --- | --- |
| *SlEIN2*  Solyc09g007870 | SET1 | F: | -1823 | 26 | CTTCGACATAATTCTCTCAAAACACC |
|  |  | R: | 468 | 24 | TCAAACACTTAAAACAGCTGGAGA |
|  | SET2 | F: | -592 | 24 | ACTGTGATGTATGTTAGGTCGTGA |
|  |  | R: | 317 | 24 | AAGAGCAGGAGAATCAACTTCG |
|  | SET3 | F: | 31 | 26 | GCTGTAGTGCTAGTTTCCTGCGATC |
|  |  | R: | 937 | 22 | ATGTCCATTCCAAATAAGTCAT |
|  | SET4 | F: | 900 | 22 | TCCAATCAAGTTACACCACTAA |
|  |  | R: | 1884 | 22 | CGTTCTCAGATACTCCTTTGAT |
|  | SET5 | F: | 1777 | 22 | TCTTCAATACTGAAACTGTGGA |
|  |  | R: | 2972 | 22 | TCTCTGCAGACTTTAGGAGAAT |
|  | SET6 | F: | 2896 | 22 | AACAAGCCTATATGAGTGGTTC |
|  |  | R: | 3996 | 26 | ATGAACTGAGAAAAGAGCGTTACAAG |
| *SlEIL1*  Solyc06g073720 | SET1 | F: | 13 | 22 | TTCCAGTTGAACCACAGTTTTG |
|  |  | R: | 1353 | 21 | CACAGTAGGTTCATTAGCTTG |
|  | SET2 | F: | 1112 | 20 | CCTTTGTCCTCAGCTGGTGT |
|  |  | R: | 2341 | 22 | AGTAAAGACCTTAACCATAGCC |
| *SlEIL2* Solyc01g009170 | SET1 | F: | 5 | 22 | CGAGTCTACCTTCTATATTCTG |
|  |  | R: | 1555 | 22 | AAGTCCAGATGTATCAAAGGAG |
|  | SET2 | F: | 1113 | 22 | GTGCTATTGATGACCCTATCTT |
|  |  | R: | 2095 | 25 | CTACTCCCTTGTAATAACGTCATGC |
| *SlEIL3*  Solyc01g096810 | SET1 | F: | 71 | 22 | CATTCTTCTCAAACCCCTTCTC |
|  |  | R: | 1224 | 22 | GTCATGGGGTTTACATTCCACT |
|  | SET2 | F: | 1077 | 22 | TTTGGCTCGTAAGCTGTATCCT |
|  |  | R: | 2148 | 22 | AGCACTTACAACCATGGAAACA |

**F**-Forward; **R**-Reverse primer

The primers were designed using the NCBI gene sequences for Ethylene Insensitive 2 (*SlEIN2,* DQ 409173), Ethylene Insensitive like 2 (*SlEIL2*, AF328785), and Ethylene Insensitive like 3(SlEIL3, AF32878) in NCBI and Solanaceae Genome Network (SGN) sequence for Ethylene Insensitive like 1 (*SlEIL1*, Solyc06g073720).
